# Screening of Chimeric GAA Variants in a Preclinical Study of Pompe Disease Results in Candidate Vector for Hematopoietic Stem Cell Gene Therapy

**DOI:** 10.1101/2021.12.28.474352

**Authors:** Yildirim Dogan, Cecilia N. Barese, Jeffrey W. Schindler, John K. Yoon, Zeenath Unnisa, Swaroopa Guda, Mary E. Jacobs, Christine Oborski, Diana L. Clarke, Axel Schambach, Richard Pfeifer, Claudia Harper, Chris Mason, Niek P. van Til

**Author notes:** To whom correspondence should be addressed (NvT); (CM). Contributed equally (Yildirim Dogan, Cecilia N. Barese, Jeffrey W. Schindler, John K. Yoon).

## Abstract

Pompe disease is a rare genetic neuromuscular disorder caused by acid alpha-glucosidase (GAA) deficiency resulting in lysosomal glycogen accumulation and progressive myopathy. Enzyme replacement therapy (ERT) is the current standard of care, which prolongs the quality of life for Pompe patients. However, ERT has limitations due to lack of enzyme penetration into the central nervous system (CNS) and skeletal muscles, immunogenicity against the recombinant enzyme, and requires life-long biweekly infusions. In a preclinical mouse model, a clinically relevant promoter to drive lentiviral vector-mediated expression of engineered GAA in autologous hematopoietic stem and progenitor cells (HSPC) was tested with nine unique human chimeric GAA coding sequences incorporating distinct peptide tags and codon-optimization iterations. Vectors including glycosylation independent lysosomal targeting (GILT) tags resulted in effective GAA enzyme delivery into key disease tissues with enhanced reduction of glycogen, myofiber and CNS vacuolation, compared to non-tagged GAA in *Gaa* knockout mice, a model of Pompe disease. Genetically modified microglial cells in brains were detected at low levels, but provided robust correction. Furthermore, an aminoacid substitution in the tag added to reduced capacity to induce insulin signaling and there was no evidence of off-target effects. This study demonstrated the therapeutic potential of lentiviral HSPC gene therapy exploiting optimized GAA tagged coding sequences to reverse Pompe disease pathology in a preclinical mouse model providing a promising vector candidate for further investigation.

**One Sentence Summary:** A candidate vector for hematopoietic stem cell gene therapy of Pompe disease.

## Introduction

Pompe disease, or glycogenosis type II (OMIM #232300), is an autosomal recessive metabolic myopathy caused by alpha-glucosidase (GAA) enzyme (EC 3.2.1.20) deficiency. It is characterized by lysosomal glycogen accumulation predominantly in the heart, skeletal muscle and CNS (1, 2). Infantile-onset Pompe disease (IOPD) patients have low level GAA enzyme activity (<2% of normal) and display severe muscle weakness. Late-onset Pompe disease (LOPD) patients have higher residual GAA activity and show a more protracted disease progression, often becoming wheelchair and ventilation-dependent, with an overall shortened life expectancy (1). The standard of care is enzyme replacement therapy (ERT), which requires lifelong weekly to biweekly recombinant human GAA (rhGAA) administrations that can prolong the life of Pompe patients but cannot guarantee long-term symptom-free survival. ERT acts through cation-independent mannose 6-phosphate receptor (CI-M6PR) also known as insulin-like growth factor 2 (IGF2) receptor (IGF2R) mediated endocytosis to the lysosome (3), which is complicated by very low rhGAA concentrations reaching the interstitial space hampering efficient uptake into affected muscle cells (4). In addition, the currently approved rhGAA has low abundance of mannose-6-phosphate (M6P), in particular biphosphorylated N-glycans, which are required for high affinity to the CI-MPR, leading to inefficient cellular uptake (4). Hence, other neo rhGAA with increased bis-mannose 6-phosphate levels have been investigated in clinical trials (avalglucosidase alfa; ClinicalTrials.gov ID: NCT02782741, NCT03019406, NCT02032524) (5) with recent FDA approval for LOPD patients. Furthermore, abnormal M6P trafficking may also hamper uptake (6, 7). To address limitations of ERT and to enhance delivery to the skeletal muscle, a GILT tag based on the IGF2 sequence that binds with high affinity to the IGF2R was fused to a truncated catalytic domain of GAA (reveglucosidase alfa) and demonstrated to be more effective in clearing glycogen in skeletal muscle in a *Gaa^-/-^* mice (3). Reveglucosidase alfa infusions in LOPD patients were reasonably well tolerated and initial improvements of respiratory strength and ventilatory function were observed, but with limited effect on walking endurance (8) (ClinicalTrials.gov ID: NCT01230801).

Alternatively, there have been efforts to treat Pompe disease patients with adeno-associated virus (AAV) gene therapy (9), as well as preclinical attempts to use the hematopoietic system as a factory to produce recombinant human (rh) GAA enzyme (10–12). For treatment of patients with lysosomal enzyme deficiencies, such as mucopolysaccharidosis I (MPSI), HSPC transplantation is a therapeutic option if a matched donor can be found. However, allogeneic bone marrow transplantation for Pompe disease patients has been unsuccessful as a treatment (13), and the low level expression of endogenous GAA enzyme in hematopoietic cells is insufficient for cross-correction. GAA activity in hematopoietic cells, such as peripheral blood (PB) leukocytes, bone marrow cells and splenocytes in mice is also low (10). Hence, allogeneic HSPC transplantation is unlikely to be beneficial and high-level vector–driven ectopic enzyme expression in hematopoietic cells would be required to accomplish efficacy. In clinical trials, lentiviral-mediated HSPC gene therapy for (neuro)metabolic diseases, such as X-linked adrenoleukodystrophy (X-ALD) and metachromatic leukodystrophy (MLD) provided therapeutic benefit and proved safe, and more recent trials initiated for Fabry disease and Hurler disease showed promising initial treatment results (14, 15). Moreover, preclinical studies using *ex vivo* LV vector-mediated overexpression of GAA using the spleen focus forming virus (SFFV) U3 promoter and partial chimerism of genetically modified cells were demonstrated to alleviate clinical symptoms in *Gaa^-/-^* mice (10). Using weaker promoters, such as locus control region of the *β*-globin chain fused to the elongation factor 1*α* promoter (LCR-EFS), that upregulate expression in the erythrocyte lineage, have been shown to be safe in murine and human HSPCs but had limited therapeutic response in skeletal muscle and CNS (16). Furthermore, in contrast to ERT and AAV therapies, in which immune responses against the rhGAA protein, viral vector or transgene product could affect efficacy (17–20), allogeneic HSPC transplantation was shown to promote immune tolerance induction to infusion of recombinant alpha-L-iduronidase (IDUA) in Hurler disease patients (21). Similarly, in preclinical studies using HSPC gene therapy in *Gaa^-/-^* mice robust immune tolerance induction, against rhGAA was also observed (10, 12).

Besides immune-related complications in AAV gene therapy, AAV serotypes targeting the muscle or liver (22) do not effectively deliver rhGAA to the CNS, and in turn, AAV serotypes targeting the CNS may not completely address the systemic pathology, such as cardiac correction (23). The more prominent CNS involvement, particularly in aging IOPD patients, could potentially be addressed through cross-correction by HSPC derived genetically modified microglia that produce the transgene product locally.

To enhance the efficacy of treatment by HSPC gene therapy, we tested ten novel lentiviral vectors expressing engineered human GILT-tagged *GAA* sequences *in vitro*. Nine of these lentiviral vectors were selected for testing in *ex vivo* HSPC gene therapy in *Gaa^-/-^* mice and provided candidates that provided biochemical correction highly efficiently in heart, skeletal muscles and CNS, showing improved efficacy compared to the codon-optimized *GAA* sequence through HSPC gene therapy in *Gaa^-/-^* mice.

## Results

### Secretion and uptake of genetically engineered GAA through IGF2R mediated pathways

Eleven different lentiviral constructs encoding unique human *GAA* sequences were generated (Fig. S1A, B). These included sequences with codon-optimized GAA (GAAco) and ten constructs containing an IGF2-tag fused to the N-terminus of the catalytic GAA sequence (GILTco) as previously described (3). Derivatives of GILTco were generated to eliminate a consensus furin-cleavage site (R37A) in the IGF2 sequence (24). Native sequence (GILTm) and three coding sequences generated through two different codon-optimization algorithms (GILTco1-m or GILT-co2-) using accession Y00839, and a third containing the consensus amino acid GAA sequence (NCBI CCDS database: CCDS32760.1 = GILT-co3-m). Another two lentiviral vectors contained a tandem repeat sequence of the apolipoprotein E (ApoE) either upstream of the GILT-tag (GILTco1-m-ApoE1) or downstream of the GILT-tag (GILTco1-m-ApoE2) to potentially facilitate enhanced blood brain barrier (BBB) crossing (25–27). Furthermore, we introduced a Gly-Ala-Pro peptide linker within the GAA amino acid sequence to enhance enzyme activity (28, 29) (GILTco1-m-L) and with the ApoE tag (GILTco1-m-ApoE1-L and GILTco2-m-ApoE2-L).

GAA is poorly secreted by cells (30). The signal peptide algorithm SignalP 5.0 (31) predicted a higher likelihood of signal peptidase Sec/SPI cleavage of the GILT-tagged GAA (0.98) compared to the untagged GAA (0.26), implying increased secretion of the GILT-tagged protein.

Lentiviral vector transduced HAP1 *GAA^-/-^* cells showed GAA enzyme activity in the cell lysates (Fig. 1A) and in conditioned media (Fig. 1B), with the GILT-tagged constructs providing 3 to 6-fold improved secretion compared to GAAco, except for the ApoE1 group, showing that GILT-tagged GAA protein is secreted more effectively than native GAA protein *in vitro*. In addition, the transduced HAP1 *GAA^-/-^* cells showed both the precursor (110 kDa) and mature protein (75-70 kDa) in the cells (Fig. S2). We further investigated cellular uptake of the chimeric proteins by IGF2R (Fig. 1C; Fig S3). GAA deficient K562 cells (K562-*GAA^-/-^*) were derived through targeted disruption of the *GAA* gene using CRISPR/Cas9 (Fig. S3A-C). Additionally, purified rhGAA, GILT-GAA and GILT-R37A-GAA protein showed similar GAA enzyme activities relative to protein concentration (Fig. S3D, E) and efficient cellular uptake by K562-*GAA^-/-^* cells (Fig. 1C). Subsequently, after disruption of *IGF2R* expression by CRISPR/Cas9 gene-editing in K562-*GAA^-/-^* cells, IGF2R protein was significantly affected as shown by flow cytometry and Western blot analysis (Fig. S3F-I), and consequently, uptake of rhGAA, rhGILT-GAA and rhGILT-R37A-GAA proteins were also significantly reduced (∼3-fold) at the highest concentrations (Fig. 1C). Reintroduction of *IGF2R* by lentiviral transduction (Fig. S3G, J) confirmed uptake through IGF2R by partially restoring cellular uptake of the three proteins (Fig. 1C). Finally, IGF2 binds to a different domain than M6P to IGF2R (3), and both GILT-GAA and GILT-R37A-GAA proteins showed specific competitive inhibition in a cellular uptake assay via IGF2 administration, but not by M6P (Fig. 1D).

**Fig. 1.**
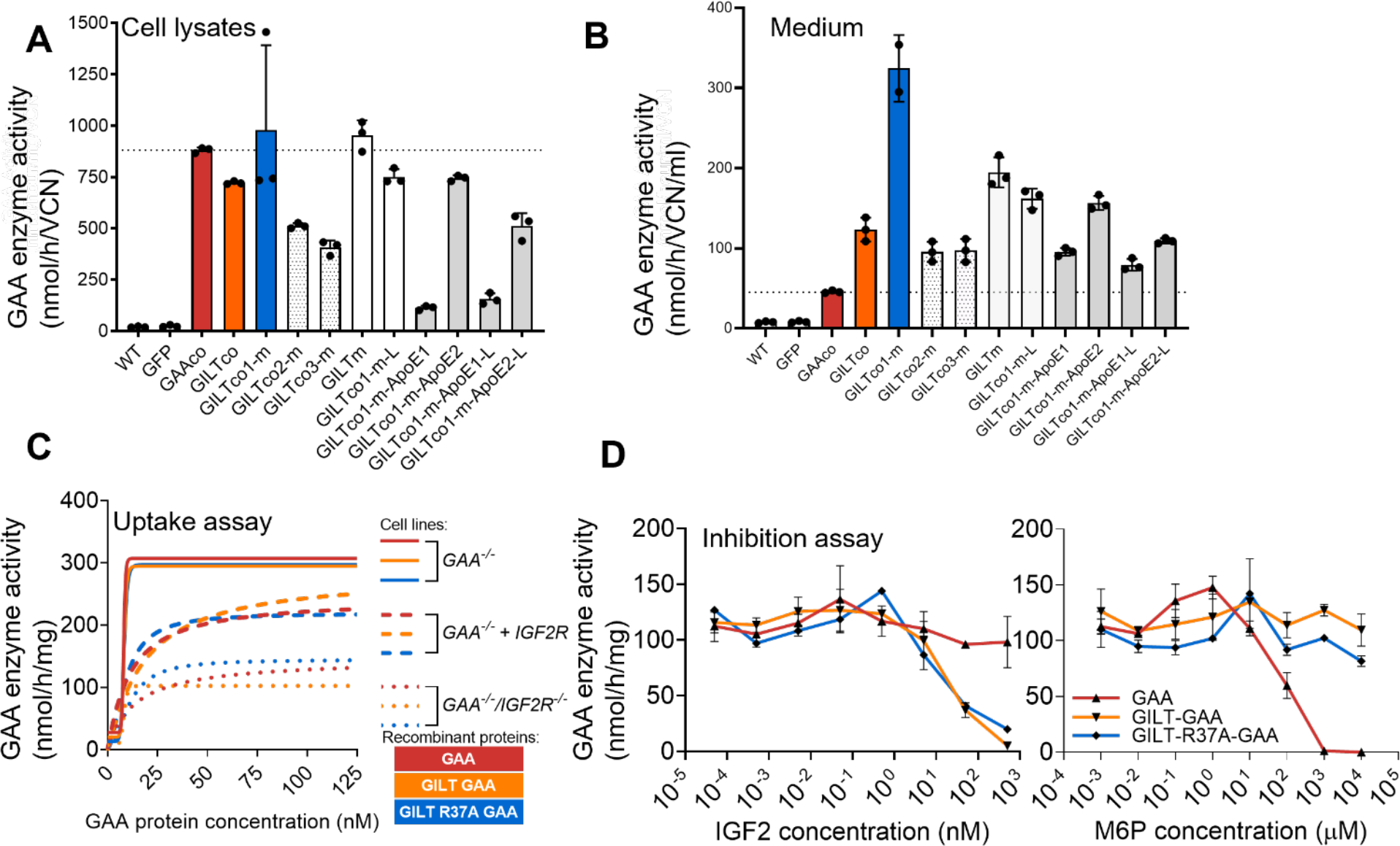
Effective secretion and uptake through IGF2R mediated pathways. (**A**) GAA activity in transduced HAP1 *GAA^-/-^* cell lysates (n=3) normalized to VCN, (**B)** and in conditioned media from the cells in Fig. 1A. (n=2-3). (**C)** GAA activity after incubation with rhGAA proteins into (1) K562-*GAA^-/-^*, (2) *GAA*^-/-^*IGF2R*^-/-^ or (3) *GAA*^-/-^*IGF2R*^-/-^ + *IGF2R* vector. Duplicates means connected by a nonlinear fit line. (**D)** GAA activity after uptake of rhGAA proteins into human Pompe patient fibroblasts. Duplicates means ± SD shown.

### Genetically modified HSPCs provide robust expression of engineered GAA in *Gaa^-/-^* mice

Donor lineage-negative (Lin-) cells derived from *Gaa^-/-^* mouse bone marrow (Fig S4A, B) transduced with the chimeric GAA vectors (nine groups) or GAAco (Table S1) were intravenously transplanted into 7.5 Gy irradiated *Gaa^-/-^* mice. The GILTco1-m variant was also transplanted in 9 Gy irradiated or Busulfex^®^-conditioned *Gaa****^-/-^*** mice. Controls included treatment naive *Gaa****^-/-^*** or *Gaa^+/+^* mice and transduction controls containing MND.GFP control groups.

Lin- cells efficiently transduced at a vector copy number (VCN) range of 0.9 to 4.6 assessed on day 7 of liquid culture (Fig. S5A). Total frequency of colony-forming units (CFU) of transduced *Gaa*^-/-^ Lin-cells showed no significant differences between non-transduced, GFP vector, or chimeric GAA vector variants (Fig S5B; p=0.277, one-way ANOVA).

The transduced *Gaa*^-/-^ Lin-cells containing chimeric *GAA* reconstituted *Gaa*^-/-^ mice and produced supraphysiological but steady increase of GAA enzyme activity in PB leukocytes ranging from ∼30 to over 100-fold increase when compared to *Gaa^+/+^* mice (median 0.64 nmol/mg/h; weeks 5; 9; 13 and 15; Fig. S5C), with GAAco, GILTco1-m-ApoE2-L, and GILTco3-m as the best performing variants. Likewise, in the GFP control groups, the percentage GFP positive PB leukocytes remained steady (mean % GFP between 45.2% ± 1.42 to 79% ± 1.38) in GFP treated mice until end of study (Fig. S5D) and high in bone marrow, and spleen (Fig. S5E). At week 16, GAA enzyme activity in the bone marrow was ∼16 to 275-fold higher in gene-therapy treated mice compared to nontreated *Gaa*^+/+^ mice (median 7.42 nmol/mg/h Fig. 2A, B) and GFP treated *Gaa^+/+^* control mice. In plasma, there was a 2.7 to 4.2-fold increase compared to the baseline signal detected in untreated *Gaa*^-/-^ and *Gaa*^+/+^ mice, and to treatment control MND.GFP groups (Fig 2 C, D). Supraphysiological GAA enzyme activities were also measured in splenocytes (Fig S5F). The GILTco1-m-ApoE1 variant showed significantly lower enzyme activity compared to GILTco and GILTco1-m (P<0.0001) in PB leukocytes, bone marrow and spleen, similar to the *in vitro* experiment. The GILTco3-m and GILTco1-m-ApoE2-L enzyme activities were significantly higher in PB leukocytes, bone marrow, spleen and plasma (P<0.05) compared to the GILTco variants. Furthermore, *Gaa^-/-^* mice treated with the GILTco1-m variant had similar GAA enzyme activity and plasma concentration across the three conditioning regimens 7.5 Gy; 9 Gy or Busulfex^®^ (Fig. 2A-D and Fig. S5C, F, H).

**Fig. 2.**
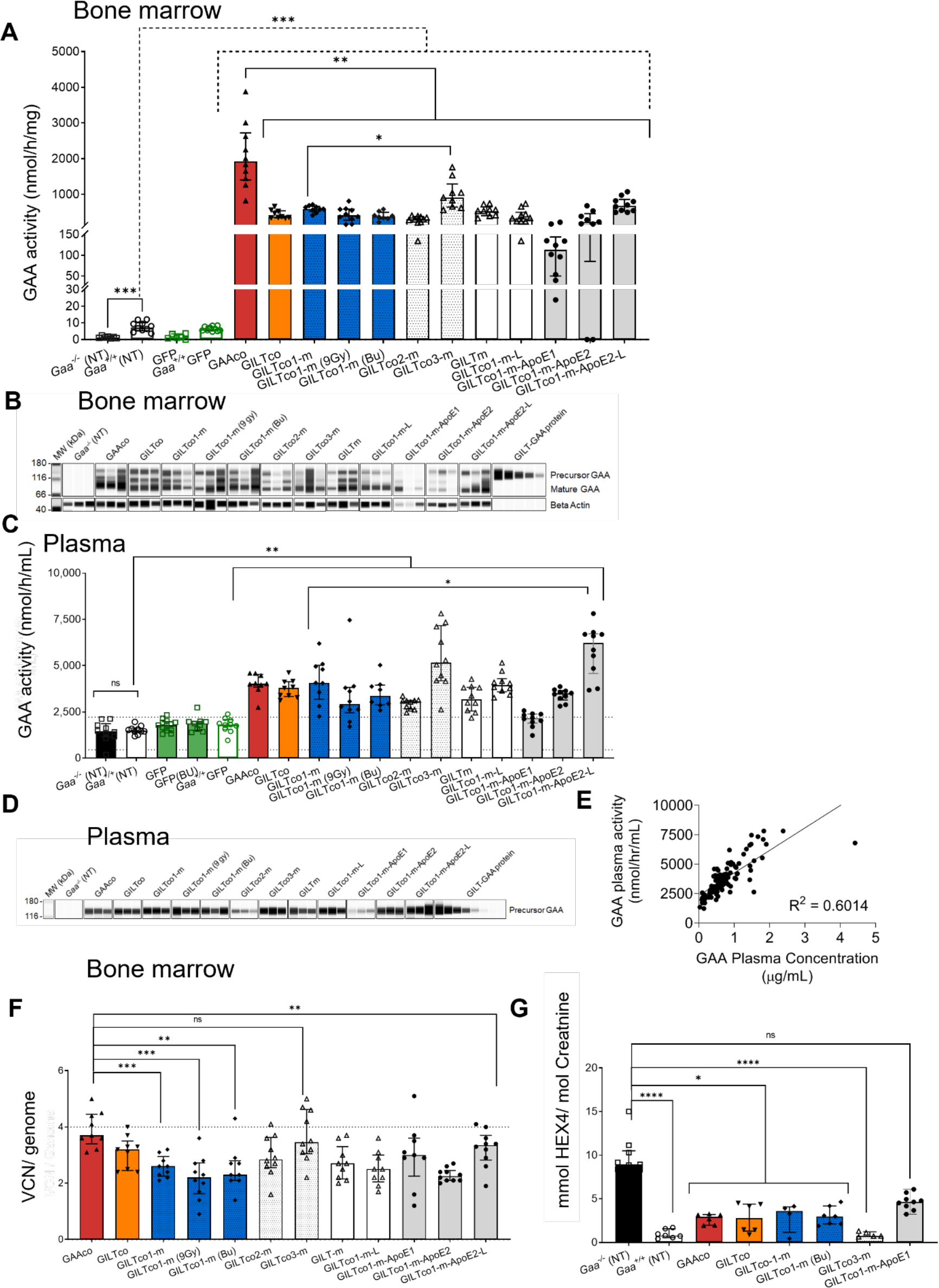
High levels of GAA in hematopoietic compartment. **(A)** GAA enzyme activity in bone marrow cells. Individual values, group medians and interquartile range presented; Comparisons: non-treated *Gaa^-/-^* vs *Gaa*^+/+^; *Gaa*^+/+^ vs treated (dotted line); GAAco vs GILT-tagged variants (solid line); GILTco1-m vs GILTco3-m (n=10). **(B)** WB analysis of bone marrow lysates. Both precursor (110kDa) and mature GAA species (75-70 kDa) was detected. Loading control: anti-beta actin (n = 3). **(C)** GAA enzyme activity in plasma at week 16 post-transplant, comparing non-treated *Gaa*^+/+^ vs treated (n=10). **(D)** WB analysis of representative plasma samples (n = 3) at week 16 post-transplant using an anti-GAA monoclonal antibody. **(E)** Correlation analysis between plasma GAA enzyme activity and plasma GAA protein concentration by simple linear regression *P*<0001. **(F)** VCN measured in bone marrow cells comparing GAAco treated group to GILT-tagged variants (n = 9-10). **(G)** Urinary HEX4 at week 16 post-transplant, individual values, group medians, and interquartile ranges presented (n=4-0). Untreated *Gaa^-/-^* mice vs GAAco Kruskal-Wallis test and Dunn’s test for multiple comparison between the treatment arms. (A, C, F) Exact Wilcoxon Rank Sum test comparing arms. *P*-values: *p< 0.05, **p<0.01, ***p<0.001 and ****p<0.0001. WB: Western blot.

Western blot analysis and quantification of bone marrow showed presence of the 110-kDa GAA precursor protein, as well as lysosomal matured 70-75 kDa GAA forms (Fig. 2B, Fig. S5G) and secreted GAA precursor protein in plasma (Fig. 2D, Fig. S5H). GAA enzyme activity in plasma ranged from 2.7 to 4.2-fold increase compared to baseline signal detected in untreated *Gaa*^-/-^ and *Gaa*^+/+^ mice, and to the treatment control MND.GFP group. A strong positive correlation was observed between GAA enzyme activity and GAA protein concentration in the plasma (Fig. 2E; P<0.0001). Quantification of the GAA protein in the plasma confirmed the highest values in groups GILTco3-m and GILTco1-m-ApoE2-L (1.66 ± 0.73, n = 20), while the lowest values were seen in the GILTco1-m-ApoE1 group (0.15 ± 0.05, n = 10). The range of all values in all groups was 0.03 to 4.4 μg/mL (Fig. S5H).

Robust transgene expression in the spleen was confirmed by *WPRE* fluorescence *in situ* hybridization (FISH) staining, and presence of GAA and the GILT-tagged protein by immunohistochemical staining in a GILTco1-m and GFP treated mouse (Fig. S5J).

Importantly, the highest median VCN in bone marrow was 3.7 for the GAAco group, but all the other therapeutic groups were lower than that (median VCN range of groups 2.2-3.4), with higher VCNs in the engrafted mice of the GFP control groups (median and range of group GFP 10.5 ± 5.8, n = 10; GFP (Bu) 9.2 ± 14.8, n = 6; and *Gaa^+/+^* GFP 9.1 ± 13.8, n = 10 ; Fig. 2F, S5K). Of note, VCNs in GAAco, GILTco3-m and GILTco1-m-ApoE2-L groups were 1.3-fold higher than in the GILTco1-m group, but GAA enzymatic activity per VCN was similar across the vector groups (Fig S5L). Furthermore, male donor cell chimerism assessed by Y-chromosome qPCR (Fig S5M) showed similar chimerism across groups and indicated that the GAA variants did not affect engraftment potential.

The urine Hex4 glucotetrasaccharide (Glc4) has been used as a biomarker to measure efficacy of ERT in Pompe patients and efficacy of AAV gene therapy in *Gaa^-/-^* mice (30, 32). In our study, Hex4 was significantly lower in the therapeutic GAAco, GILTco, GILTco1-m, GILTco3-m groups compared to the *Gaa^-/-^* mice at 16 weeks (Fig. 2G), suggesting improved glycogen clearance in skeletal muscles.

### Genetically modified HSPCs with engineered GAA clear glycogen in heart and skeletal muscles

At 16 weeks post infusion of genetically modified HSPCs, GAA activity in the cardiac tissue in the majority of gene therapy treated *Gaa^-/-^* mice was restored to normal levels as in the wildtype *Gaa*^+/+^ group and *Gaa*^+/+^ GFP mice (6.3-6.4 nmol/h/mg; Fig. 3A), with the exception of the ApoE-tag-containing GILTco1-m-ApoE1 variant and the GILTco1-m-ApoE2 variant (± 2.7 fold lower than the GILTco1-m group; p<0.0001, and p<0.0164, respectively). The highest GAA enzyme activity was observed in the GAAco group (21.5 nmol/mg/h), followed by GILTco-3-m group (14.7 nmol/mg/h), with 2.3-fold excess of wildtype *Gaa^+/+^* mice (Fig. 3A). Of note, GAA enzyme activity in the GILTco1-m group (7.5 Gy) was similar to 9 Gy and Busulfex^®^ mice. Heart GAA enzyme activities aligned with observed GAA protein concentration (Fig. 3B and S6A), and mature lysosomal GAA protein (70-75kDa) was most prominently detected (Fig. 3B). Previous studies showed that lentiviral HSPC gene therapy using a *GAA* transgene effectively reduced heart glycogen levels (10, 11, 16), but also in our study more than 99% reduction was observed in all the therapeutic groups (except GILTco1-m-ApoE1) at week 16 (Fig. 3C) when compared to *Gaa^-/-^* mice (441.44 μg glycogen/mg protein; Fig. 3C). Quantification of Periodic Acid Schiff (PAS) and hematoxylin and eosin (H&E) staining performed in hearts of selected groups (Fig. 3D; Tables S2,3 and Fig. S7,8) confirmed significant reduction of glycogen and typical myofiber/vascular vacuolation in the GAAco group, but even more so in the GILTco, GILTco1-m, and GILTco3-m groups. Furthermore, cardiac hypertrophy is a hallmark of IOPD patients and *Gaa^-/-^* mice (1, 33). Importantly, significant heart mass reduction was observed in the non-GILT-tagged and GILT-tagged groups acquiring similar heart mass as the *Gaa^+/+^* group, except in the GILTco1-m-ApoE1 and GILTco1-m-L groups, which was not statistically significant from *Gaa^-/-^* mice (Fig. 3E).

**Fig. 3.**
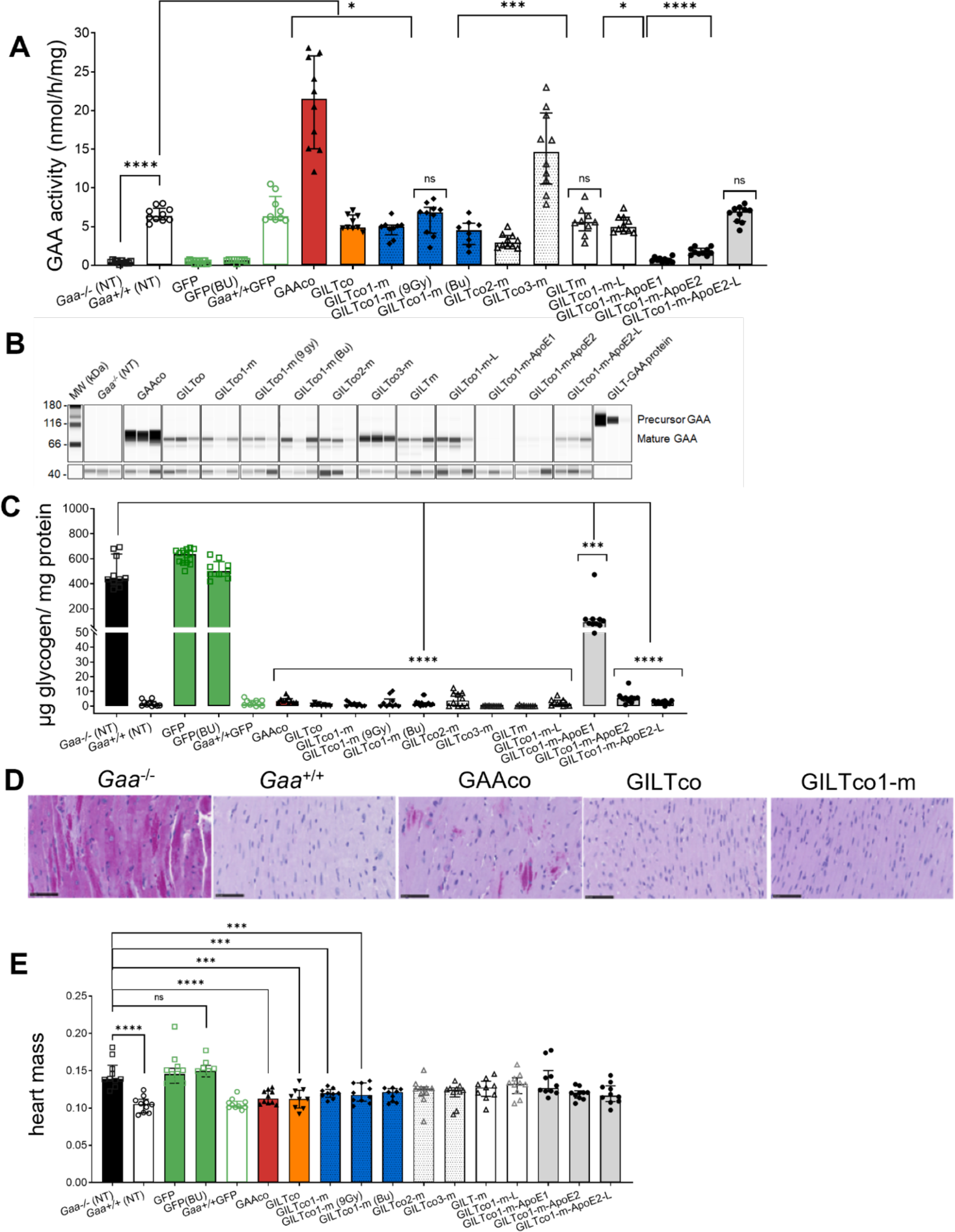
Robust correction in heart of *Gaa^-/-^* mice. **(A)** GAA enzyme activity in the heart of treated *Gaa^-/-^* mice; *Gaa^+/+^* vs treatment groups. (**B)** WB analysis of heart tissue lysates with presence of mature GAA species (75-70 kDa; n = 3). (**C**) Reduction of lysosomal glycogen in the heart is shown with treated compared to non-treated *Gaa^-/-^* mice. (**D**) Representative PAS staining images of heart for glycogen (purple stain) of non-treated vs treated groups. (Scale bar = 50µm). (**E**) Reduction of heart mass with group comparison between *Gaa^-/-^* vs *Gaa*^+/+^ and vs GAA variants is shown (n = 8-10). Individual values, group medians and interquartile ranges are shown. Wilcoxon Rank Sum test p value comparing arm vs *Gaa*^+/+^ or *Gaa*^-/-^. *P*-values: **p<0.01, ***p<0.001 and ****p<0.0001.

In the diaphragm and the hindleg muscles quadriceps, tibialis anterior, and gastrocnemius, the highest GAA enzyme activity was detected in the GAAco group followed by the GILTco3-m group (average all skeletal muscles: 5.7-fold and 3.5-fold increase to *Gaa^+/+^* respectively; Fig. 4A-D, left). In the majority of other therapeutic groups, GAA enzyme activity was restored to normal levels in *Gaa^+/+^* mice. Western blot analysis of gastrocnemius tissue (Fig. 4E and S6B) affirmed presence of more GAA protein (70-75 kDa) in the GAAco and GILTco3-m group than all other groups. Using lentiviral HSPC gene therapy with a *GAA* transgene, skeletal muscle, such as diaphragm, but more prominently hindleg muscles such as quadriceps, tibialis anterior, and gastrocnemius muscles which were historically more resistant to glycogen clearance (10), in our study showed robust glycogen clearance across all groups (Fig. 3A-D, right), except the GILTco1-m-ApoE1. Of note, in the skeletal muscles glycogen was significantly more reduced in the GILT-tagged groups, particularly the GILTco1-m (∼3% in diaphragm and 24 – 35% in hindleg muscles of *Gaa^-/-^* (NT)) and GILTco3-groups (∼1% in diaphragm and 10 – 17% in hindleg muscles of *Gaa^-/-^* (NT)), which were ∼2.3-fold and ∼5-fold lower, respectively, compared to the GAAco group (P-value <0.001; glycogen in GAAco group in diaphragm = 12%, gastrocnemius = 77%, quadriceps = 57%, and tibialis = 60% of *Gaa^-/-^* (NT) group). PAS quantification (Fig. 4F, S7 and Table S2) in selected samples confirmed the greater reduction in glycogen and myofiber vacuolation pathology in the GILTco1-m and GILTco3-m groups (Fig. S8 and Table S3).

**Fig. 4.**
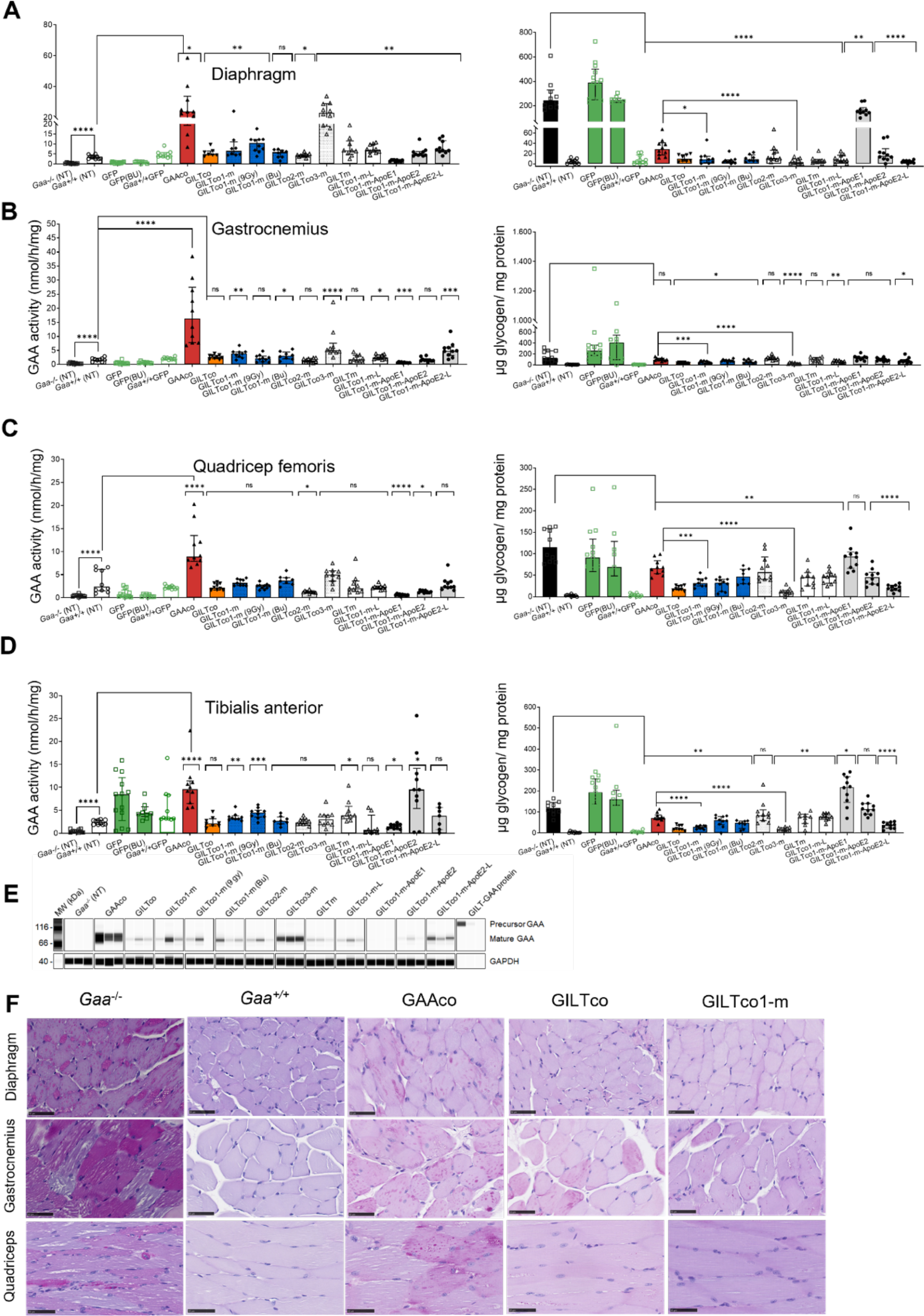
Effective correction in skeletal muscles of *Gaa^-/-^* mice by GILT-tag. Left panel shows GAA activity, right panel shows glycogen content. (**A)** diaphragm (**B)** gastrocnemius (**C)** quadricep femoris (**D)** tibialis anterior. Individual values, group medians and interquartile ranges. Exact Wilcoxon Rank Sum P-value comparing with *Gaa^+/+^* or *Gaa^-/-^* group. *P*-values: *p< 0.05, **p<0.01, ***p<0.001 and ****p<0.0001 (n = 8-10). (E) WB analysis of representative gastrocnemius tissue lysates with presence of mature GAA species (75-70 kDa). Loading control: GAPDH. (n = 3). (F) PAS stainings of heart, diaphragm and quadriceps for glycogen (purple stain) of treated vs non-treated samples (Scale bar =50µm).

### GILT-R37A-tag rescues central nervous system pathology

We then investigated whether HSPC gene therapy could penetrate the CNS and provide biochemical correction through local presence of genetically modified microglia. GAA enzyme activity in the cerebellum in the GAAco, GILTco, GILTco-1-m (9 Gy), GILTco1-m-ApoE2, and GILTco1-m-ApoE2-L groups were significantly higher compared to untreated *Gaa^-/-^* mice (p < 0.0001; p 0.0279; p 0.0029; p 0.0029; p 0.0288; respectively) (Fig. 5A, left). However, the GAA enzyme activity levels never came close to nontreated *Gaa^+/+^* mice, e.g. wildtype levels of ∼30% with GAAco, 18% with GILTco1-m (9 Gy), and less than ∼6% in the other therapeutic groups. In contrast, in the cerebrum only the GAAco group showed detectable GAA activity that was above non-specific signal in *Gaa^-/-^* controls (1.16 nmol/mg/h vs 0.36 nmol/mg/h; p 0.0001, GAAco vs *Gaa^-/-^*, respectively) (Fig.5B, left). Mature GAA protein was predominantly observed in cerebrum, which was ∼6.7-fold higher in the GAAco group when compared to the GILT-tagged groups (Fig 5C, Fig S6C).

**Fig. 5.**
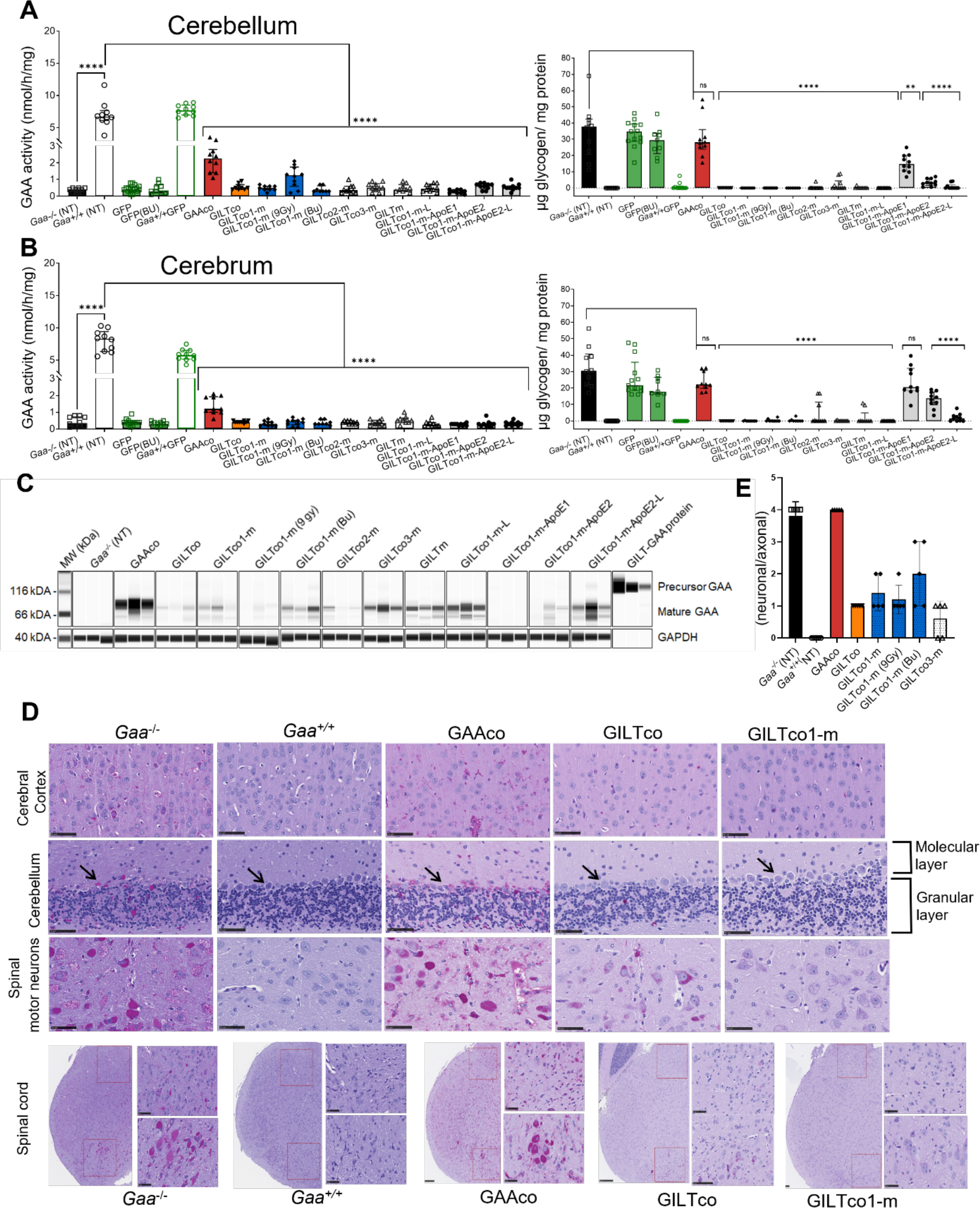
GILT-tag GAA required for resolving pathology in CNS. GAA enzyme activity in the cerebrum and cerebellum of *Gaa^-/-^* mice. (**A, B**) Left: GAA activity; Right: Glycogen. Values are for individual animals, represented as group medians and interquartile range. Exact Wilcoxon Rank Sum p-value comparing treatment groups vs *Gaa* ^+/+^ or *Gaa^-/-^* (n= 8-10)*. P*-value *p< 0.05, **p<0.01, ***p<0.001 and ****p<0.0001. **(C**) WB analysis of cerebrum detecting mature GAA species (75-70 kDa). Loading control: GAPDH (n = 3). (**D)** PAS staining images of brain and spinal cord sections for glycogen with glycogen (purple stain) depicted in non-treated vs treated samples. Top: Cerebral cortex, cerebellum and spinal motor neurons. Cerebellum sections show the molecular and granular layers, arrow indicates Purkinje cells (Scale bar=50µm). Bottom: PAS staining of the cervical spinal cord regions, indicating the dorsal horn region (top right panel) and the ventral horn region (bottom right panel; scale bar =250µm for high and 50µm low magnification). (**E)** Neuronal/axonal vacuolation severity scores.

Despite the low GAA activity and protein content in cerebellum and cerebrum, glycogen was significantly reduced (e.g. <99% in GILTco-1-m, and GILTco-3-m), except in the GAAco (p = ns, both 74% vs *Gaa^-/-^* (NT) groups) and GILTco1-m-ApoE1 (p ns) groups, which were indistinguishable from *Gaa^-/-^* mice (Fig.5A, B right), with the GILT-tagged variants showing superior performance, which was maintained with the R37A substitution. Furthermore, reduction in PAS staining intensity of the cerebral cortex, cerebellum, and cervical spinal cord was observed, including in spinal cord motor neurons (Fig 5D, for quantification see Fig S7 and Table S2). As a consequence of glycogen reduction, neuronal/axonal vacuolation was reduced in the GILT-tagged groups (Fig. 5E). In addition, vascular vacuolation ranged from minimal to moderate in the *Gaa^-/-^* controls, however, in GILT-tagged treated mice vascular vacuolation was minimal to mild (Table S4).

To assess conditioning to promote engraftment of donor genetically modified cells in the brain of *Gaa^-/-^* mice, we utilized the GFP vector groups. We found that the percentage of GFP positive cells in the total brain that co-expressed the microglial marker Iba1 was 51.2% ± 17.6% (n= 21) across all groups, and that cells that showed co-localization exhibited a microglial morphology (Fig 6A). The extent of genetically modified microglia in the brain was on average across all groups for total brain 1.7% ± 1.06% (n = 21, Fig. 6B) as assessed by Iba1+/GFP+ positive cells, comparable to cortex, and hippocampus. Furthermore, co-localization of Iba1+ and *WPRE*-containing transcripts by FISH in multiple brain regions of GILTco1-m treated mice showed rare positive cells in multiple brain regions, although the signal was highly variable between animals (Fig 6C), e.g. average across the three 7.5 Gy, 9 Gy and Busulfex^®^ treated GILTco1-m in cortex 0.6% ± 0.9 (n=21), and in hippocampus 1.8% ± 3 (n=21; Fig 6D). Total levels of Iba1+ cells in the cerebral cortex and hippocampus showed no significant difference between control *Gaa*^-/-^, *Gaa*^+/+^, or the .5 Gy, 9 Gy or Busulfex^®^ GILTco1-m groups (Fig S9A).

**Fig. 6:**
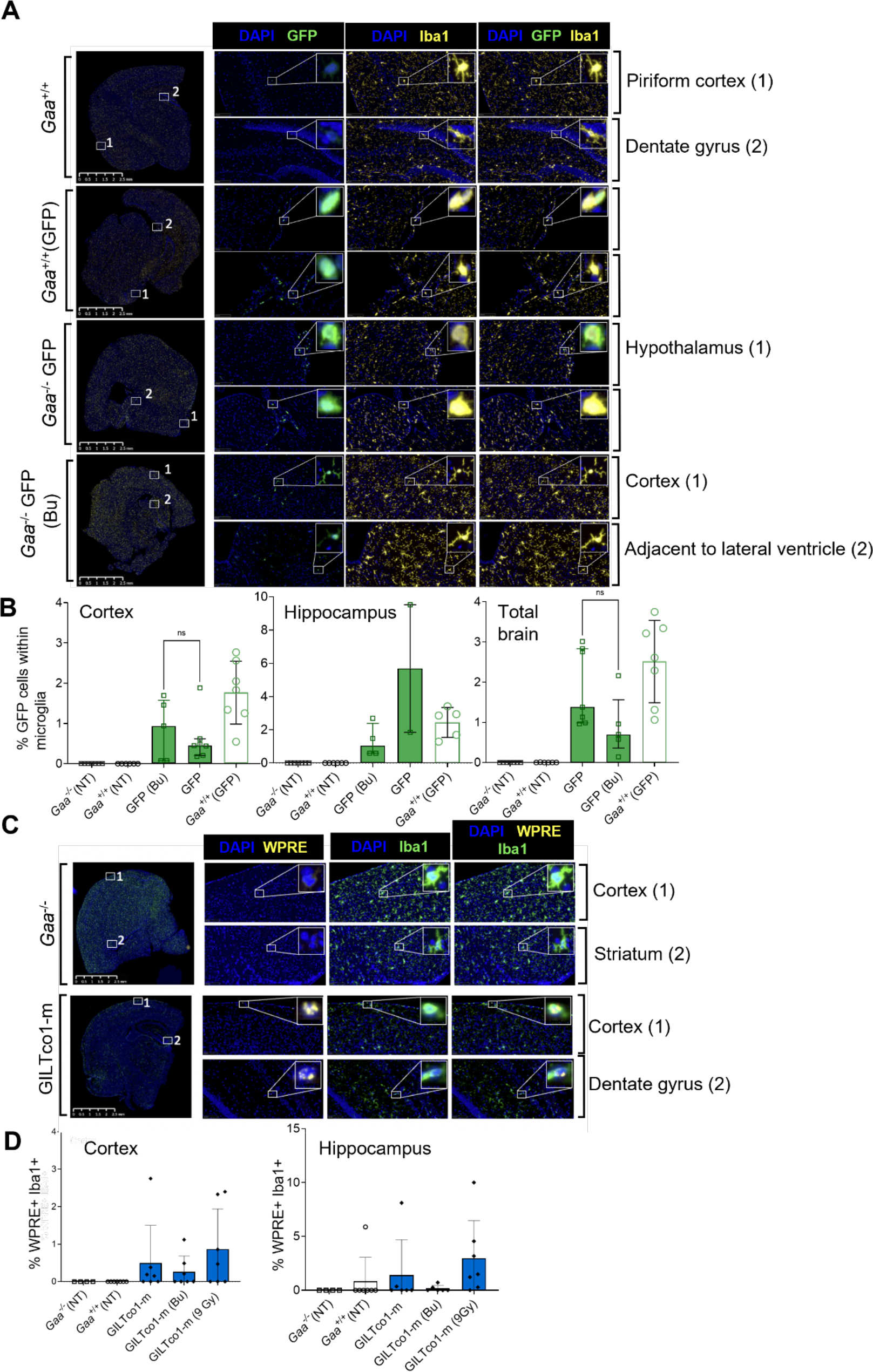
Lentiviral vector expression in brain structures. **(A)** GFP-positive microglia in brain tissues. Representative images of regions of interest (ROI) showing Iba1 positive microglia (yellow) with amoeboid to ramified morphology (higher magnification). ROI included piriform cortex dentate gyrus, hypothalamic nucleus, adjacent to lateral ventricle, hypothalamus, reticular nucleus, and cortex. *Gaa*^+/+^ animals served as controls. (**B)** % GFP+ cells in the microglia (Iba1+) cells in the cortex, hippocampus, and total brain (n=5-7). (**C)** FISH colocalization of *WPRE* RNA (yellow) signal within the microglial cells (Iba1+, green immunofluorescence) in one gene therapy treated group. Top: *Gaa*^-/-^ animals Bottom: GILTco1-m. Representative ROIs include cortex (all groups), striatum (*Gaa^-/-^*) and dentate gyrus (GILTco1-m). (**D)** %WPRE+ cells in the Iba1+ microglial cell population for the cortex and hippocampus (n = 4-7).

### GILT-R37A does not perturb insulin signaling, blood glucose or hematopoiesis

The use of reveglucosidase alfa, an IGF2-tagged GAA analog, was able to induce transient hypoglycemia at high dose infusions in LOPD patients (8). Hence, we developed a reporter assay to investigate the ability of the GILT-R37A-tag modification to induce insulin signaling (Fig S10). High rhGILT-R37A-GAA protein concentrations did not produce a reporter signal, as compared to strong responses detected by low concentrations of insulin, and 10-1000-fold lower activation by IGF2 and GILT-GAAco addition (Fig 7A). Additionally, if off-target effects such as insulin signaling would have occurred by application of the GILT-R37A-tag, we would expect a reduction in glucose levels in treated mice as well as increased mortality. Nevertheless, blood glucose levels were similar across treatment groups and controls until the end of study (Fig. 7B glycemia prior termination, week 4, 8, 12 not shown) and there was no treatment-related mortality in this study. Other potential long-term blood glucose perturbations, such as assessed by glycated hemoglobin levels (% HbA1c), were within normal range in therapeutic and controls groups, suggesting no obvious evidence of blood glucose dysregulation in the GILT-tagged GAA treated mice (Fig. 7C). Finally, hematopoiesis was assessed in PB at monthly intervals, and at termination. The erythroid, platelet as well as myeloid and lymphoid populations, i.e. CD11b+Gr1+ myeloid cells, B220+ B lymphocytes, CD3e+CD4+ and CD3e+CD8+ T lymphocytes populations were comparable between the different study groups and conditioning regimens over time (Fig. S10, Table S5, week 16). At termination, bone marrow and spleen populations were similar across groups (Table S6, S7). Overall, this demonstrated that lentiviral HSPC gene therapy using GILT-R37A or other tag combinations did not affect engraftment and differentiation in *Gaa*^-/-^ recipient mice. The supraphysiological GAA levels were well tolerated with no increased mortality. Only one animal each of the GILT-m, GILTco1-m, GFP and GILTco groups were found dead or preterminally euthanized, but these were isolated events unrelated to the vector constructs tested.

**Fig. 7.**
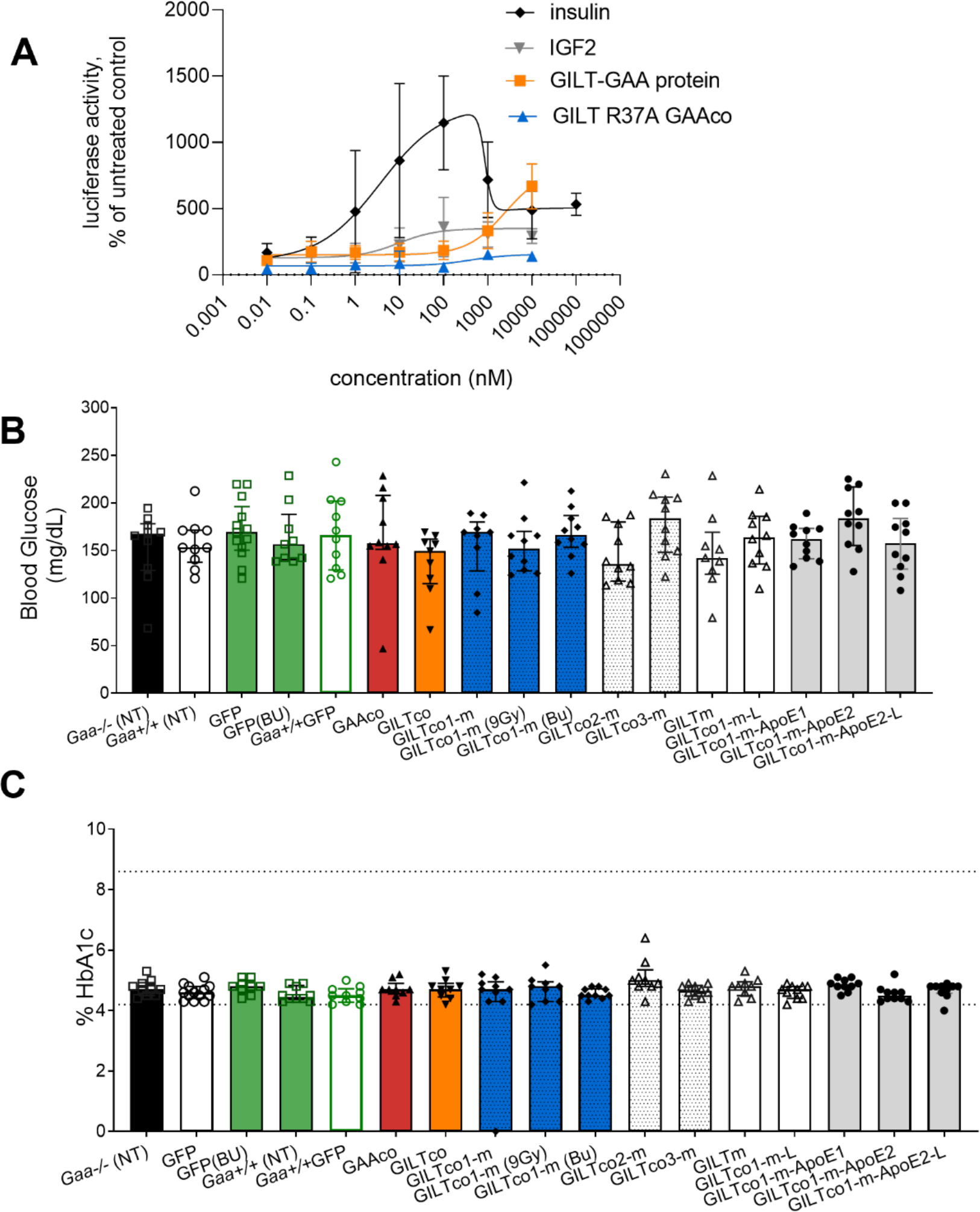
GILT-R37A does not evoke an insulin signaling response. **(A)** RAT2 fibroblast insulin reporter cells were treated with 0.01nM -100µM concentrations of insulin, IGF2, rhGILT-GAA or rhGILT-R37A-GAA protein. Each data point presented as mean (triplicate) ± SD. Results were normalized to untreated cells. (**B)** Glycemia (**C**) and HbA1c blood levels in experimental mice prior termination, groups medians with interquartile ranges (dot lines show normal ranges; n = 9-13).

## Discussion

In this study, we describe nine lentiviral vectors using a clinically relevant promoter driving expression of engineered GILT-tagged GAA variants for improved efficacy in HSPC-mediated gene therapy. Most of these GAA variants conferred increased secretion from hematopoietic cells, and biochemical correction and alleviation of pathology in key target tissues including heart, skeletal muscle and CNS. The conditions tested in this study surpassed previous results of preclinical studies using lentiviral HSPC gene therapy, in which a lentiviral vector with SFFV promoter was used to express *GAA* (10) or a codon-optimized GAA (11).

The SFFV promoter has been used in clinical gene therapy trials using long terminal repeat (LTR)-driven gammaretroviral vectors for chronic granulomatous disease (CGD) (34, 35) and adenosine deaminase (ADA) severe combined immunodeficiency (SCID) gene therapy (41). The field is moving away from the gammaretroviral vector HSPC gene therapy due to genotoxicity risks, and recent trials have adopted third generation self-inactivating lentiviral vectors with more physiological promoters with improved efficacy and safety (36, 37). In our approach, the MND promoter was incorporated in vector design. MND is a promoter that has been used for gammaretroviral vector ADA-SCID HSPC gene therapy with a follow-up protocol of more than 8-11 years, in which nine out of ten patients had functional immune system recovery with no need for ERT or a secondary allotransplantation. To our knowledge, no adverse events related to this gammaretroviral vector were noted (38). Moreover, the MND promoter has also been used for lentiviral HSPC gene therapy in cerebral adrenoleukodystrophy (CALD) with reported follow-ups of four patients up to 12 years after transplantation (14) and another study with a shorter follow-up of 17 CALD treated patients (39). In addition to these ongoing studies, another HSPC gene therapy trial for treatment of recombination activating gene 1 (*RAG1*) severe combined immunodeficiency (SCID) children adopted a vector design using the MND promoter to drive RAG1 expression, and this study is open for patient recruitment (ClinicalTrials.gov ID: NCT04797260). Hence, to provide robust expression of our chimeric *GAA* coding sequences, the MND promoter was incorporated in our third-generation self-inactivating lentiviral vector backbone that is currently applied in a follow-up phase 1/2 lentiviral HSPC gene therapy for Fabry patients (40) (ClinicalTrials.gov ID: NCT04999059) and first-in-human study for Gaucher patients (41) (ClinicalTrials.gov ID: NCT04145037).

In other lysosomal disorders, such as Hurler and MLD, endogenous expression of the lysosomal protein has been shown insufficient to provide cross-correction (42) emphasizing the need to gain therapeutic potential by boosting therapeutic protein levels. The current standard of care for Pompe disease requires high dose infusions of rhGAA protein (alglucosidase alfa; 20 mg/kg bodyweight, biweekly), which is approximately 13-20-fold higher than recombinant protein therapies in other LSDs such as Fabry disease (agalsidase beta 1 mg/kg bodyweight) (43) or Gaucher disease (1-1.5mg/kg bodyweight) (44). More recently, even higher (40mg/kg) and more frequent doses (weekly) are infused to benefit Pompe patients (17, 45), but the approved standard of care is limited by the uptake in target cells, poor receptor recycling on target tissue, and its inability to cross the BBB. A rhIGF2-tagged GAA, which was designed to improve muscle uptake, was shown to increase uptake in L6 myoblasts 25-fold and was significantly more effective in reducing glycogen in multiple skeletal muscles compared to rhGAA (3). Using GILT-tag technology, the requirement of high abundance biphosphorylated glycans for sufficient uptake in target tissues could be circumvented. In another AAV-mediated gene therapy study, replacing the endogenous GAA signal peptide with a signal peptide of a secreted protein enhanced excretion from hepatocytes, thereby improving cross-correction to target tissues (22, 30). Most chimeric GAA proteins tested in our study were also effectively produced from cells and secreted into conditioned media. In addition, we showed that the GILT-R37A-GAA protein lacking the consensus furin cleavage site in the IGF2 moiety was effectively taken up through its IGF2 binding site on IGF2R in cellular assays.

Altogether, our results show robust expression using GILT-tagged *GAA* transgenes compared to the native GAA sequence and broad biodistribution of the GAA proteins to relevant tissues, demonstrating an efficacious advantage that was most prominently apparent in the CNS. The GILT-R37A-tag conveyed an enhancement to the safety profile of this approach, by considerably reducing the affinity of the protein to the insulin receptor and showed similar efficacy. In many studies, optimization of transgene expression and protein translation via the recoding of the transgene has been successful for HSPC lentiviral gene therapy with up to 10-fold higher transgene expression, e.g. recombination activating gene 1 (*RAG1*), and 2 (*RAG2*), common gamma chain (*IL2RG*) and beta-glucocerebrosidase (*GBA*) (46–49). However, the codon-optimization algorithms investigated in our study did not improve rhGILT-R37A-GAA protein production, similar to another study using codon-optimized *GAA* for AAV therapy (28). Hence, the *GAA* sequence may already be an optimal sequence for mammalian transcription and translation.

Tagging protein technology has been employed to deliver gene-therapeutics to the brain in preclinical studies using IGF2 moieties to target the CI-M6P receptor through intrathecal delivery in MPSIIIB mice (50), targeting the transferrin receptor through fusing antibody domains to therapeutic proteins, such as progranulin for neurological disorders (51) or systemic delivery using ApoE tandem tags in MPSII mice (27). However, incorporating additional CNS-specific tags, such as ApoE, in an attempt to enhance BBB-crossing delivery, did not enhance CNS pharmacodynamics in our study (27). In our case, the ApoE tag was fused to the N-terminus upstream or downstream of the IGF2 moiety and divided by a short GAP linker. In the study of Gleitz *et al*. a larger flexible peptide linker was used to separate the ApoE-tag from iduronate-2-sulfatase enzyme (IDS) and the tag was fused to the C-terminal end, which resulted in efficient reduction of brain heparan sulfate compared to the non-tagged IDS protein (27). In addition to that, as Gleitz *et al* hypothesized, the ApoE moiety may have provided a stabilization effect to the IDS protein in plasma, which was not apparent in our study. The context-dependent positioning of IGF2, ApoE and GAA peptide sequences may have affected the effective production, at least for the ApoE1 construct. It may also have hampered the capacity of the ApoE-tagged products to dock to their cognate receptors the low-density lipoprotein receptor (LDLR) or LDLR-related protein 1 (LRP1) (26) from target cells or interfere with IGF2R receptor binding, although this would require further investigation. Finally, a GAP linker in the protease 3 site (29), which was reported to improve enzyme activity after transient transfection (28) did not provide enhanced biochemical correction in our study.

*In vivo* AAV gene therapy has been investigated in preclinical models in *Gaa^-/-^* mice using different routes of administration mainly using AAV1, AAV8 and AAV9 serotypes targeting liver, muscle or CNS (58) and a clinical trial directing AAV1 serotype to the diaphragm (9). However, long-term efficacy can be significantly hampered by the complexity of immune responses to the transgene products or CD8+ T cell immune responses to the AAV vector capsid (52). The use of AAV vectors and restricting expression to hepatocytes could induce immune tolerance induction against rhGAA (53). A major potential advantage of HSPC gene therapy is immune tolerance induction, which has previously been demonstrated in *Gaa^-/-^* mice (61, 62). Since HSPCs are transduced *ex vivo*, immune responses to vector components are not anticipated. Our study was a 16-week follow-up of hematopoietic reconstitution of genetically modified Lin-bone marrow cells, in which GAA enzyme activity in PB and plasma was stable for all therapeutic treatment groups and no indication was seen of an immune response to the recombinant proteins.

Heart and diaphragm tissue typically responds more efficiently than skeletal muscle to treatment in *Gaa^-/-^* mice and IOPD patients (54, 55). In our study, all chimeric constructs except the ApoE1-tagged variant reduced glycogen significantly in heart (<99%). However, when using other non-systemic gene therapy approaches, such as intrathecal delivery of AAV9 in *Gaa^-/-^* mice, it was more challenging to correct the cardiac phenotype (23). In another study, a secretable GAA was effective in reducing the glycogen in both the heart and skeletal muscle (22). Using the GILT-tagged GAA constructs, the skeletal muscles diaphragm and gastrocnemius responded well with near complete glycogen reduction, but quadriceps and tibialis anterior were more resistant and contained more residual glycogen. The best results were obtained with the GILTco3-m, confirmed by histological analysis, most likely because tissue enzyme activities were higher than the other groups, which can be explained by the modestly higher VCNs measured *in vivo*. In addition to the reduction of glycogen, a reversal of myofiber and vascular pathology in the skeletal muscle was also observed, with vacuolation severity and incidence being decreased in all animals treated with GILT-tagged GAA constructs tested, compared to animals treated with the non-tagged construct. It is becoming more apparent that ERT cannot address CNS involvement, which is a key component in the Pompe disease phenotype (56). The BBB limits the effectiveness of large molecule therapeutics, and is also often compromised by lysosomal disease (57). Confounding effective delivery is the requirement of M6P receptors for uptake and lysosomal trafficking, which are expressed at low levels during adulthood (58). As an example, neither imiglucerase nor velaglucerase alfa passes the blood-brain barrier in Gaucher patients. For this reason, ERT is not seen as efficacious for Gaucher patients with the acute neuronopathic form (type 2), and consequently there is no substantial change in the life-threatening neurological parameters (59). Similarly, rhGAA does not cross the BBB effectively, as has been observed in long-term ERT-treated LOPD patient after autopsy, who had significant motor-neuron glycogen depositions in the ventral horn of the spinal cord (60). The use of lentiviral HSPC gene therapy to deliver the non-tagged *GAA* transgene product to the CNS was not effective in reducing lysosomal glycogen (10). In another approach, using liver-directed AAV8 vectors producing chimeric GAA to enhance secretion and reduce transgene product immunity of *Gaa^-/-^* mice, the CNS was also not effectively treated (22). In another study, intralingual injection of AAV9 to deliver IGF2-tagged GAA tongue myofibers and motor neurons in *Gaa^-/-^* mice resulted in a >200% increase in the number of GAA-positive motor neurons compared to untagged GAA with clearance of glycogen (61), showing increased effectivity of an IGF2 tag approach for CNS delivery. Interestingly, in our study, although GAA enzyme activity was detectable in the brain of mice in the GAAco group at approximately 11% and 30% of wildtype levels in cerebrum and cerebellum respectively, glycogen level was not reduced. This confirms previously reported results, in which glycogen in the CNS was not reduced through HSPC gene therapy using solely the *GAA* sequence (10, 16). Surprisingly, although GAA enzyme activity in the GILT-tagged groups in the brain was below detectable enzyme activity levels, lysosomal glycogen was completely reduced to wildtype levels, which was confirmed by near absence of histological PAS staining. These histopathological findings in the CNS of GILT-tag treated *Gaa^-/-^* mice showed reduced glycogen accumulation in both neurons and glial cells in the cerebral cortex, corpus callosum, hippocampus and cerebellum. In brainstem and spinal cord, PAS staining revealed extensive glycogen reduction in the large neurons, especially in the motoneurons. In the brain stem, cerebellum, cerebral cortex and spinal cord, PAS staining approached wild type levels in all GILT-tagged treated groups, but not in the non-tagged GAAco treated group. Western blot protein quantification on cerebral homogenates across groups, excluding the ApoE-tagged variant groups, showed average GILT-tagged protein levels between 5 – 30% when compared to the GAAco group, which demonstrates that even low levels of GILT-tagged enzyme was sufficient in significantly reducing glycogen and vacuolization.

We hypothesize that the mechanism of the transgene product delivery to the brain could potentially go through two processes: transcytosis of GAA protein through the BBB or through local production of HSPC derived microglia in the brain. Microglia constitute 5-12% of rodent brain (62) transplanted and non-transplanted *Gaa^-/-^* mice in our study had a range of 13.3%-23% Iba1+ cells in the cortex, and a range of 7.9%-12.8% Iba1+ cells in the hippocampus across groups. In a study, in which irradiation and the two alkylating agents treosulfan and busulfan were tested, conditioning was instrumental for microglia-derived engraftment in the brain, but busulfan was not favorable over the other treatments (63). However, in another study, in which busulfan and irradiation conditioning were compared, busulfan enhanced donor-derived engraftment in the brain (64). In our study, we followed engraftment in the GFP treated groups and examined colocalization with Iba1+ staining for microglia and noted that 0.1-3.7% of the total Iba1+ cells were GFP+, with no evident differences between 7.5 Gy, 9 Gy or Busulfex^®^ conditioned groups. Based on these percentages, the percentage of genetically modified microglia of the total population in the brain is therefore estimated to be around 1%, which explains why the GAA enzyme activity was below detectable levels. Presence of genetically modified therapeutic Iba1+ cells in the brain was confirmed by *WPRE* FISH, and therefore local delivery of GILT-tagged GAA could explain the lysosomal glycogen reduction observed. It is remarkable that the low enzyme activity levels were sufficient to reduce overall glycogen in the brain in *Gaa^-/-^* mice. Similar low enzyme activity levels were observed in other studies using HSPC gene therapy; e.g. in *Arsa^-/-^* mice, not more than 10% of wildtype arylsulfatase A enzyme activities were detected in brains of HSPC lentiviral vector *ARSA* treated mice (65), and in *Idua^-/-^* mice lentiviral HSPC gene therapy resulted in 4.5-fold wildtype levels (66). In another LSD rodent model, treated *Ids^-/-^* mice reached enzyme levels that did not exceed 4% of wildtype levels in brain, but heparan sulfate and dermatan sulfate levels were completely cleared (27). Similarly, the results from our study show that GAA enzyme activity and transgene product measured in the CNS, whether it is derived from native GAA or GILT-tagged GAA, does not directly correlate with the reduction of glycogen. Nonetheless, it is promising that very low levels of brain chimerism were adequately mitigating brain pathology. Effective substrate breakdown has been shown using IGF2-tagged NAGLU under conditions of limited exposure on MPSIIIB patient fibroblasts, mouse primary astrocytes, and cortical neurons, by which heparan sulfate was more effectively broken down, and showing the improved uptake by these specific cell types (50). This enhanced uptake may aid in the reduction of glycogen in the CNS of *Gaa^-/-^* mice. We do not rule out that transcytosis, in part, also plays a role in the delivery of GILT-tagged protein to the CNS, but this would require further investigation. If local production of the transgene product would be the sole mechanism to reduce pathology in the brain, further enhancement of efficacy may be achieved by supplementing FDA-approved compounds, such as G-CSF1R inhibitors to enhance engraftment of HSC-derived microglia in the CNS (67).

In a clinical trial infusing 20mg/kg reveglucosidase alfa in LOPD patients (mean weight 89.0 kg and range 49.2 – 144.5kg) hypoglycemia was observed (8), because IGF2 binds to the insulin receptor (IR), and particularly with higher affinity to the IR absence (IR-A) variant that lacks exon 11 (68). Hence, we generated a cell line containing *IR-A*, *STAT5b* and containing a *STAT5*-luciferase reporter based on a previously published reporter cell line to assess insulin receptor kinase activity (69). Using this insulin receptor reporter assay, insulin signaling by GILT.R37A.GAA protein could not be detected, confirming the enhanced safety of the R37A mutein.

Furthermore, low levels of between 0.03 – 4.4 μg/mL of GILT-tagged GAA protein were measured in plasma in our gene therapy treated mice. Based on an adult blood volume of 5 liters and dose of 20 mg/kg, the concentrations that we detected in our gene therapy-treated *Gaa^-/-^* mice were estimated to be at least 45-131-fold lower than the concentrations estimated in the plasma of LOPD patients treated with IGF2-tagged GAA who experienced hypoglycemia adverse events (8). The highest GAA protein concentration measured in our gene therapy treated mice was 4.4 μg/mL, but the average protein concentrations were actually *∼* 3 – 7-fold lower than that. We postulate that these low concentrations of circulating GILT-tagged GAA are expected to significantly reduce the likelihood of hypoglycemia events. This line of reasoning is supported by the observation that there were no reported hypoglycemia events in the ERT clinical trial treatment group receiving a low dose of 5 mg/kg (8). Of note, in the reveglucosidase alfa clinical trial, none of the subjects in the study required dose modification or discontinued treatment because of hypoglycemia (8). No hypoglycemia was observed in our study; glucose levels were in the range of untreated *Gaa^-/-^* mice, and average glucose levels as assessed by hemoglobin A1c levels were also not perturbed.

Altogether, based on our findings, a lentiviral vector containing IGF2-tag with R37A substitution (GILTco1-m) was selected as a promising candidate for assessment in long-term efficacy, biodistribution and toxicology preclinical evaluation and potential application in first-in-human lentiviral HSPC gene therapy of Pompe disease.

## Materials and Methods

### Study Overview

To evaluate the therapeutic effect of lentiviral HSPC gene therapy for the treatment of Pompe disease, a lentiviral vector containing codon-optimized *GAA* and constructs containing IGF2 and/or ApoE moieties, linker inclusions and different codon-optimization sequences were generated. These configurations were tested in HAP1-*GAA^-/-^* cells for protein production and secretion and K562 cell lines were generated to assess uptake by the GILT-tag. A reporter cell line was also generated to assess insulin signaling by the IGF2 and IGF2-R37A moieties. After *in vitro* assessment, nine GILT-tagged *GAA* containing lentiviral vectors were selected for *in vivo* testing in female B6;129-Gaatm1Rabn/J [6neo] mice (*Gaa* knockout; *Gaa^-/-^*) mice. All animal experiments were approved by the local Institutional Animal Care and Use Committee. After conditioning the mice with 7.Gy or 9 Gy gamma-irradiation or Busulfex^®^ (4 x 25/mg/kg), 6-9 weeks old mice were injected intravenously with enriched male donor Lin-HSPCs transduced with engineered *GAA* vectors encoding for native or chimeric GAA proteins. Female *Gaa^-/-^* and *Gaa^+/+^* mice GFP vector and nontransduced control groups were included. Mice were monitored for 16 weeks with interim blood collections. Therapeutic endpoints included biochemical GAA enzyme activity, tissue glycogen content, Western blotting analysis, VCN analysis, histological evaluation, blood glucose, glycated HbA1c, clinical pathology, immunophenotyping via flow cytometry and immunofluorescence and FISH analysis of gene modified cells in brain sections. Experimental groups were sized to allow for statistical analysis; not all the animals were included in the analysis, and select outliers were excluded. Mice were assigned randomly to the experimental groups based on weights.

### Plasmid construction and lentiviral vector production

Human codon optimized *GAA* (*GAA*co), and GILT-tagged chimeric *GAA* coding sequence variants GILTco, GILTco1-m, GILTco-m-ApoE1, GILTco-m-ApoE2, GILTco1-m-L, GILTco1-m-ApoE1-L and GILTco1-m-ApoE2-L were designed using GenSmart™ Codon Optimization Tool (GenScript). Human *GAA* coding sequence variant GILTco2-m was designed using GeneArt codon optimization algorithm (Thermofisher Scientific). The human *GAA* coding sequence variant GILTco3-m was designed using a consensus sequence (CCDS32760.1). GILT-m contained the native nucleotide sequences. All *GAA* coding sequence variants were synthesized and cloned into a lentiviral vector backbone described previously (41). The EFS promoter from this construct was replaced by the myeloproliferative sarcoma virus enhancer, negative control region deleted, dl587rev primer-binding site substituted (MND) promoter (70). Plasmids were fully verified by Sanger sequencing and produced at Aldevron.

All third-generation self-inactivating LV vectors described above were produced by transient transfection of 293T cells using packaging and transfer plasmids at the Viral Vector Core of Cincinnati Children’s Hospital. HOS cells used to determine vector titers were acquired from ATCC (Cat# CRL-1543). Titers determined on HOS cells by qPCR were in the range of 2.25 *×*10^7^ to 6.67 *×*10^7^ transducing units per mL and were subsequently titrated on lineage negative bone marrow cells to determine the multiplicity of infection (MOI) needed to achieve a VCN of ∼4.

### *In vitro* experiments for secretion and uptake of chimeric proteins

HAP1 *GAA*^-/-^ cells (Horizon Discovery, ID: HZGHC004204c004, and parental ID C631) were transduced with eleven lentiviral vectors encoding GAA variants (Fig. S1B) and GFP at an MOI of 3. Conditioned media of vector transduced cells and cell pellets were collected at day 11 for VCN normalized GAA enzyme activity measurement.

Recombinant human GILT-R37A-GAA protein was produced in HD Chinese hamster ovary cells and purified (Genscript). Cell culture supernatant was centrifuged, followed by filtration, and loaded onto affinity purification column. Eluted fractions were pooled, concentrated, and loaded onto gel filtration chromatography column. Purified protein was analyzed by SDS-PAGE, Western blot analysis, and Bradford assay to determine concentration, molecular weight and purity.

K562 cells (ATCC), K562 cells *GAA^-/-^* clone 20, *GAA^-/-^IGF2R^-/-^* clone 25, *GAA^-/-^IGF2R^-/-^* clone 25 + lentiviral vector IGF2R (Clone 51) were used for uptake assays. All knockout cell lines were generated by CRISPR/Cas9 gene-editing using ribonucleoprotein complexes (Figure S3). Knockout efficiency was assessed with surveyor assay, single cells sorted, expanded and screened for IGF2R protein by anti-IGF2R antibody (Biolegend) by flow cytometry.

K562 cell lines were incubated with concentrations of 125, 31.3, 7.8 and 2.0 nM purified rhGAA, rhGILT-GAA and rhGILT-R37A-GAA proteins at 37 °C at 5% CO2 for 18 hours. Subsequently, cells were pelleted, washed three times with DPBS, and lysed with 100µL PC-T lysis buffer. After removal of debris by centrifugation at 10,000 g for 10 min, duplicate lysate samples were used to measure GAA activity with 4-methylumbelliferyl (4MU) normalized to protein concentration with bicinchoninic acid protein (BCA) assay (Pierce).

Healthy and Pompe-affected human fibroblast cells were obtained through Coriell Institute (cat# GM07525 and GM00244, respectively). Fibroblast cells were seeded and grown for 24 h to approximately 90% confluency. Cell media was replaced with medium containing 50 nM of purified rhGAA, rhGILT-GAA and rhGILT-R37A-GAA proteins and incubated at 37°C at 5% CO2 for 18 hours. Select wells also contained inhibitors M6P (1 nM – 10 mM; Calbiochem) or IGF2 (5e-2 pM – 5e2 nM; Cell Sciences). Subsequently, cells were trypsinized, pelleted and washed three times with DPBS, and ran in a GAA activity assay.

### Insulin reporter assay

RAT2 fibroblast cells (ATCC) were genetically modified with a lentiviral vector to express human insulin receptor A (IR-A) isoform, and signal transducer and activator of transcription 5B (STAT5b) and mCherry (VectorBuilder). Cells were sorted for IR-A and mCherry, and subsequently transduced with a puromycin-selectable STAT5 response element luciferase reporter lentiviral vector (G&P Bioscience). Following puromycin selection, the RAT2 reporter cell line was plated at a concentration of 100,000 cells/well in a 96 well plate in DMEM, 10% FBS, 100U/ml penicillin and 100µg/ml streptomycin. After 12 hours, media was replaced, supplemented with 0.01nM-100µM concentrations of insulin, IGF2 or GILT-GAA or GILT-R37A-GAA protein. Following 8 hours of activation with the supplemented media, One-Glo Luciferase Assay reagent was added to the wells. Following 10 minutes of lysis, lysates were transferred to a white walled 96 well assay plate for measurement on the Softmax i3x plate reader. Results were normalized by dividing each treated well values by the mean of the untreated values.

### Mice

The *Gaa^tm1Rabn^*/J mice (*Gaa^-/-^* mice, Pompe mice) were used for the study (33). Mice were obtained from an on-site breeding colony (kindly provided by Dr. Nina Raben, NIH, MD). Control B6129SF1/J mice (*Gaa*^+/+^ mice, WT mice) were obtained from Jackson Laboratories (Stock No. 101043). All mice were maintained in clean rooms and fed with irradiated certified commercial chow and sterile acidified water *ad libitum*. Assessment of animal health status, body weight, and examinations during the in-life term of the study were conducted by veterinarian personnel and documented. All protocols were approved by the Institutional Animal Use and Care Committee at the Canadian Council on Animal Care.

### Assessment of dosing formulations and infusion in conditioned *Gaa^-/-^* mice

Recipient female *Gaa^-/-^* mice (6-9-week-old) were sublethally irradiated once with a gammasource at 1.6 Gy/min with a total of 7.5 Gy or 9 Gy. Another group received four intraperitoneal injections of 25 mg/kg Busulfex^®^ (Otsuka Pharmaceutical) from days -4 to -1 before cell dosing.

Bone marrow cells were harvested from femurs and tibias of 6-12-week-old male *Gaa^-/-^* donor mice, and Lin-enriched for HPSCs using RoboSep^TM^ (StemCell Technologies). After enrichment, cells were overnight transduced with ten lentiviral vectors (Fig. S1), at a density of 10^6^ cells/mL (MOI of 4) in serum free StemMACS^TM^ medium containing 100 ng/mL recombinant murine stem cell factor (SCF); 50 ng/mL recombinant human FMS-like tyrosine kinase 3 ligand (Flt-3) and 10 ng/mL recombinant human thrombopoietin (TPO; StemCell Technologies). On the following day, cells were washed, counted, assessed for viability. A 3x10^3^ cell fraction of non-transduced and transduced Lin-cells were transferred into 3 ml methylcellulose-based medium with mouse cytokines optimized for hematopoietic progenitor cell differentiation (StemCell Technologies) and plated at 1 mL/well for a CFU assay, CFUs were scored using STEMvision (Stemcell Technologies) at day 7.

Preconditioned female *Gaa^-/-^* recipients were injected intravenously with 0.5x10^6^ cells per mouse (Table S2). At four interim time points after cell infusion, leukocyte and plasma GAA enzyme activity, blood glucose, and complete blood counts using a Hemavet (Drew Scientific) analyzer were measured.

### Analysis of glucotetrasaccharide HEX4 in urine samples via LC-MS/MS

Mice were fasted before urine collection for glucotetrasaccharide HEX4. The quantification of 6-*α*-D-glucopyranosyl maltotriose (HEX4) was conducted in a Shimatzu (Columbia, MD) via protein precipitation using acetonitrile and 4 mM uric acid with 0.2% NH4OH as surrogate matrix.

Calibration standards and quality controls were run in surrogate matrix and mouse urine matrix in duplicate. Results were normalized by measurement of creatinine in the urine sample via a commercial kit.

### Blood glucose and glycated hemoglobin monitoring

Blood glucose was measured at different time points during the in-life study and prior to scheduled termination with glucometer Accu-chek AVIVA. Animals were fasted overnight prior to collections and results reported as nmol/L (standard Canadian units) and converted to mg/dL by multiplying by a factor of 8. Similarly, glycated hemoglobin (HbA1c) was measured using A1cNow (pts Diagnostics, Indianapolis, IN) and results reported in %HbA1c. Quality controls of A1cNow kit were performed with NOVA-ONE kit levels 1 & 2 (NOVA-ONE Diagnostics, Calabazas, CA).

### Flow cytometry

Peripheral blood, bone marrow, thymus and spleen cells underwent red blood cell lysis and were stained with monoclonal antibodies against surface markers CD45.1, CD45.2, CD3e, CD4, CD8a, B220, CD11b, Gr-1 (Ly6G and Ly6C), TER119, and CD41, or Sca-1c-Kit, and a hematopoietic lineage cocktail (BD Biosciences, Biolegend) and analyzed on a FACS Lyric flow cytometer using Fixable Viability Dye eFluor® 506(FVD506) (Invitrogen) for dead cell exclusion.

### Scheduled termination procedure

Scheduled termination was week 16 post-transplant. Animals were fasted 8-12 hours prior to euthanasia and anesthetized with isoflurane. After cardiac puncture blood collection, the animals were perfused with 1X PBS pH 7.4 until internal organs were pale in appearance. The heart mass was weighed, and the dissected tissues were snap frozen and stored at -80°C for GAA enzyme activity and glycogen measurement or processed for histopathology.

### Measurement of GAA enzyme activity and glycogen by biochemical analysis

For GAA enzyme activity and glycogen measurement, tissue samples of heart, diaphragm, gastrocnemius, quadriceps femoris, tibialis anterior, cerebellum, cerebrum were collected at necropsy were processed chilled to homogenization in sterile dH2O, centrifugation, removal of clear supernatants, and then stored at -80°C until the assay day. The GAA enzyme activity was measured following Jack *et al*. (71). Results were normalized for protein concentration in the sample using BCA kit (Pierce, ThermoScientific). The glycogen in tissues was estimated treating samples with and without *Aspergillus niger* amyloglucosidase following the procedure described by Okumiya *et al* (72).

### VCN analysis and donor cell chimerism

Lin-cell dosing formulations cultured until day 7 and bone marrow samples collected at termination were processed for gDNA and quantified with Quant-iT™ assay kit or Nanodrop One. This qPCR assay consisted of oligonucleotide primers and probe mix containing either a TaqMan 6-carboxyfluorescenin (FAM) or VIC^TM^ fluorescent probe designed to amplify *HIV Psi* vector and *Gtdc1* housekeeping gene sequences. A plasmid containing both sequences was used as a reference standard in a range of 50 to 5e10^7^ copies. Data was reported as VCN/diploid genome.

Comparative male donor cell chimerism was determined by PCR on bone marrow gDNA using Y chromosome gene *Zfy1* and *Bcl2* gene to normalize the total gDNA input per reaction following An et al (73). Donor-derived male cells engraftment in female recipient mice was calculated using a gDNA standard curve with known percentage of male vs. female gDNA.

### Western blot (WB) analysis

Monoclonal antibodies against GAA and GILT-tag sequence were generated by Genscript. Mouse plasma samples were screened for the presence of GAA protein using Jess^TM^ ProteinSimple and following manufacturer’s protocol (ProteinSimple, Bio-Techne Brand). The sensitivity of the method is in the pg order. Samples were denatured at 95°C before analysis. For tissues, roughly 200 ng of cell lysate was loaded per well, calculated based on the results of a BCA kit (Pierce, ThermoScientific). Antibody clone 1C12C11.F9 was used as detection antibody for GAA and clone 7C10 was used as detection antibody for the GILT tag. Dilutions of rhGAA BioMarin protein (BMN701) were included either as a positive control (tissues) or standard curve (plasma) for all assay runs. Beta-actin was visualized via ab8224 (Abcam) and GAPDH via 2118L (Cell Signaling Technologies). All Jess detection was performed as single-plex reactions in the chemiluminescent channel. Relative GAA quantification of tissues was calculated using the loading control and reported as a percentage in reference to GAAco. GAA absolute quantification in the plasma was determined by interpolating values using a BMN701 standard curve, multiplying by the dilution factor and adjusted per mL of plasma.

### Periodic Acid-Schiff (PAS) and H&E staining of mouse tissues

Following scheduled necropsy, sections cerebral cortex, cerebellum, hippocampus, and/or brainstem, thoracic and cervical spinal cord, heart, quadriceps femoris, diaphragm, gastrocnemius, and tibialis anterior processed for Periodic Acid-Schiff and hematoxylin and eosin (H&E) staining, and evaluated for glycogen accumulation and vacuolation by light microscopy. For severity scoring of vacuolation, score was assigned as minimally (score 1) affected tissues having < 50% of cells within the section with small discrete centralized regions of cytoplasmic vacuolation involving < 10% of the cytoplasmic volume; mildly (score 2) affected tissues having larger regions of vacuolation involving ≥ 10% of the cytoplasmic volume affecting < 50% of cells within the section and none to rare myofibers that were diffusely enlarged with overall decreased cytoplasmic staining intensity; moderately (score 3) affected tissues having regions of cytoplasmic vacuolation involving >10% of cell with > 50% of myofibers showing evidence of myofiber degeneration characterized by enlargement of myofibers and overall decreased staining intensity; and markedly (score 4) affected tissues having overall enlargement and decreased staining intensity of the majority of myofibers with both centralized regions of cytoplasmic vacuolation and evidence of myofiber degeneration.

### Immunofluorescence and vector RNAscope imaging of brain

The brain tissue sections were processed into three levels (forebrain, midbrain and hindbrain) following necropsy. Spleen and brain tissues were formalin-fixed and paraffin-embedded blocks were used for immunofluorescence detection of GFP with anti GFP antibody (ab290 Abcam) or rabbit monoclonal anti-Iba1 antibody (ab178846, Abcam). For the detection of lentiviral vector sequences within the tissue an untranslated mRNA sequence of Woodchuck hepatitis virus posttranscriptional regulatory element (*WPRE*) was used as a probe in fluorescent RNAscope^®^ *in situ* hybridization assay. The detection was validated by using Fluorescent RNAscope^®^ VS Universal HRP Assay. The images were scanned on a Hamamatsu NanoZoomer whole-slide scanner. Brain tissues were scanned at 40x magnification, and scans were imported into the Visiopharm software system. Regions of interest were drawn for various brain structures and each ROI was manually refined to exclude meninges and histological artifacts.

### Statistical analysis

GAA enzyme activity, glycogen, Hex4 measurement were analyzed by VERISTAT, Inc applying median and inter-quartile range using Kruskal-Wallis and exact Wilcoxon rank-sum tests for groups comparison and SAS/STAT^®^ software v9.4. A result of < 0.05 was indicative of significant difference in the groups. For correlation analysis, the Pearson R coefficient and p-value was used. Quantitative immunofluorescent data were collected in Visiopharm and statistical analysis was performed using SAS software (v9.4). Continuous variables were analyzed via Levene’s test for equality of variance. Where Levene’s test was not significant, one-way ANOVA was used to detect differences among three more group means. Tukey’s test was used to explore pairwise group comparisons when one-way ANOVA was significant. Where Levene’s test was significant, the Kruskal-Wallis test was used to detect differences between three or more group means. The Dwass-Steel-Critchlow-Fligner method was used to explore pairwise group comparisons when Kruskal-Wallis test was significant. Significance was set to p < 0.05 for all statistical tests. In the figures *P<0.05, **P<0.01, ***P<0.001 and ****P<0.0001.

## List of Supplementary Materials

### Materials and Methods

Lentiviral constructs and *in vitro* assays

Fig. S1 Graphical depiction of the lentiviral vector proviral vector and GAA sequences.

Fig. S2 Western blot analysis of HAP cell lysates.

Fig. S3 Generation of K 752 562 GAA and IGF2R KO cells.

Fig. S4 Illustration of experimental enrichment and transplant model.

Fig. S5 Characterization of lineage-negative enriched bone marrow cells and recipient hematopoietic tissues and plasma.

Fig. S6 Western blot GAA protein relative quantification percentages.

Fig. S7 PAS quantification.

Fig. S8 Myofiber vacuolation quantification.

Fig. S9 Quantification of total Iba+ cells in multiple brain regions.

Fig. S10 Generation of insulin reporter cell line.

Fig. S11 Complete blood count results in transplanted mice at week 16.

Table S1. Study design of the Vector Comparison Study (CRL #2018-3835).

Table S2. PAS quantification in CNS and skeletal muscle.

Table S3. Vacuolation scores in heart and skeletal muscle.

Table S4. Vacuolation and degeneration scores in CNS.

Table S5. Flow cytometry immunophenotyping of the peripheral blood at week 16 post-transplant.

Table S6. Flow cytometry immunophenotyping of the bone marrow and spleen at week 16 post transplant.

Table S7. Flow cytometry immunophenotyping of the spleen at week 16 post-transplant.

## Supporting information

Suppl_vanTil_et_al

## Acknowledgments

We thank all the members of AVROBIO, Inc. for their continued support of our work. In particular, we would like to thank Maurine Braun, Bianling Liu, Robert Plasschaert, Mark DeAndrade, Claudia Fiorini, Vicky Chen, Tara Peterson, Daniella Pizzurro, Nicole Del Signore, Haydy George Leha, Maria Grigorova, Steven Tyler, Becky Reese, Carolina Romano, for scientific support and assistance. We would like to also thank Jon Lebowitz (BioMarin Pharmaceutical, Inc) for providing recombinant human IGF2-GAA (BMN 701) and valuable input for the project. We would like to thank Charles River Laboratories, including Maia Araujo Abrahim, Karen Wong, Romain Genard, Lauriane Padet, Anne Larrivée, Camille Laure-Pittet, Simon Authier and Daphne Gordon for study execution and diligent care of study animals. Additionally, we thank Li Na, Kyle Takayama and Lyn Wancket (Charles River Laboratories), Jennifer Dannehl, Leonard Jared, Rathna Veeramachaneni, and Dawn Dufield, (KCAS Bioanalytical & Biomarker Services) for their contributions to sample analysis, and Douglas Arbetter (VERISTAT, Inc) for biostatistical analysis. Finally, we would like to thank Nina Raben (National Institutes of Health, MD) for supplying the *Gaa* knockout mice.

## Author contributions

YD, CB, JY, JS contributed to the experimental design and execution, biochemical and molecular assays development and qualification, data analysis and interpretation, and wrote the manuscript; ZU, SG, MJ conducted experiments, and performed data analysis, DLC organized production of lentiviral vectors, AS contributed to vector, cDNA and experimental design; CO, RP, CH contributed to assay development and study logistics; CM contributed to the design and the reporting of all work presented, NvT designed the study plan, vector configurations, performed data interpretation, and wrote the manuscript.

