## Supplementary material for "Screening of Chimeric GAA Variants in a Preclinical Study of Pompe Disease Results in Candidate Vector for Hematopoietic Stem Cell Gene Therapy": Suppl_vanTil_et_al

Materials and Methods

Fig. S1 Graphical depiction of the lentiviral vector-mediated proviral and GAA sequences.

Fig. S2 Western blot analysis of GAA and GILT-tag in HAP cell lysates.

Fig. S3 Characterization of gene-edited K562 GAA and IGF2R knockout cells.

Fig. S4 Illustration of experimental enrichment and transplant model.

Fig. S5 Characterization of genetically modified cultured donor cells and recipient hematopoietic compartment.

Fig. S6 Relative GAA protein levels in heart, gastrocnemius and cerebrum in treatment groups.

Fig. S7 Quantification of PAS staining in heart, skeletal muscles and CNS.

Fig. S8 Quantification of myofiber vacuolation in heart and skeletal muscles.

Fig. S9 Quantification of total Iba+ cells in multiple brain regions.

Fig. S10 Diagram presenting the generation of insulin reporter cell line.

Fig. S11 Complete blood count results in transplanted mice.

Table S1. Study Design of the Vector Comparison Study.

Table S2. PAS+ quantification in CNS and skeletal muscle.

Table S3. Vacuolation scores in heart and skeletal muscle.

Table S4. Vacuolation and degeneration scores in CNS.

Table S5. Flow cytometry immunophenotyping of the peripheral blood at week 16 post-transplant.

Table S6. Flow cytometry immunophenotyping of the bone marrow at week 16 post-transplant.

Table S7. Flow cytometry immunophenotyping of the spleen at week 16 post-transplant.

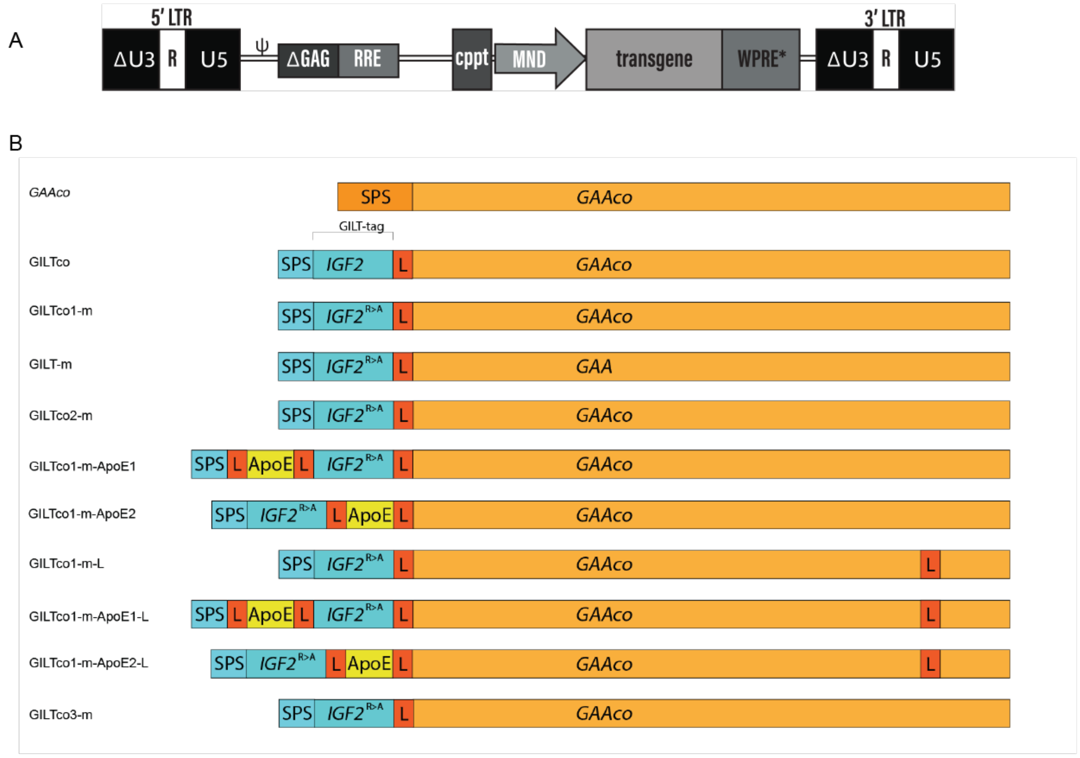


**Fig. S1: Graphical depiction of the lentiviral vector-mediated proviral and GAA sequences**.

(**A**) Schematic of proviral vector: 5' LTR = HIV 5' long terminal repeat with inactivated (delta) Unique 3', Repeat, Unique 5'; ψ = packaging signal; delta GAG = truncated HIV GAG sequence; RRE = Rev-Response Element; cPPT = central polypurine tract; MND = MND promoter; Transgene, cDNA of interest listed under Fig. S1B, WPRE* = mutated Woodchuck hepatitis virus posttranscriptional regulatory element, and 3' LTR= inactivated (delta) Unique 3', Repeat, Unique 5'. (**B**) Overview of modified acid alpha-glucosidase (*GAA*) sequences. IGF2 = insulin-like growth factor 2; SPS = signal peptide sequence (orange, native GAA SPS and blue, IGF2 SPS); 'co' = codon optimized; GILT = glycosylation independent lysosomal targeting; 'L' = Gly-Ala-Pro peptide linker; R>A = Arginine to Alanine substitution; ApoE = Apolipoprotein E tag. Note, cDNA sequences were codon optimized using Genscript's or Thermofisher's GeneArt algorithm.

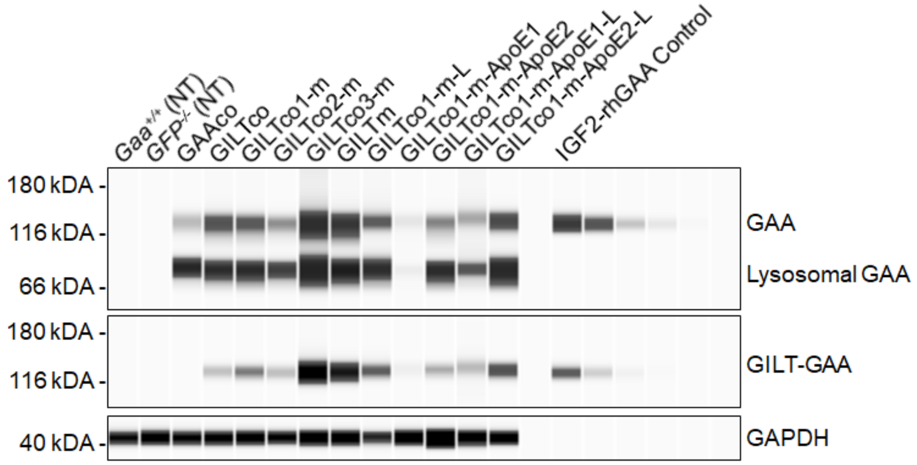


**Fig. S2**. **Western blot analysis of GAA and GILT-tag in HAP1 cell lysates.**

Top: WB analysis of transduced HAP cell lysates (MOI 3) with all *GAA* containing vectors, using an anti-GAA monoclonal antibody. Middle: WB analysis of transduced HAP cell lysates with all *GAA* containing vectors, using an anti-GILT monoclonal antibody. An anti-GAPDH antibody was used for loading control. All three blots were run using the same cell lysate master mix preparation.

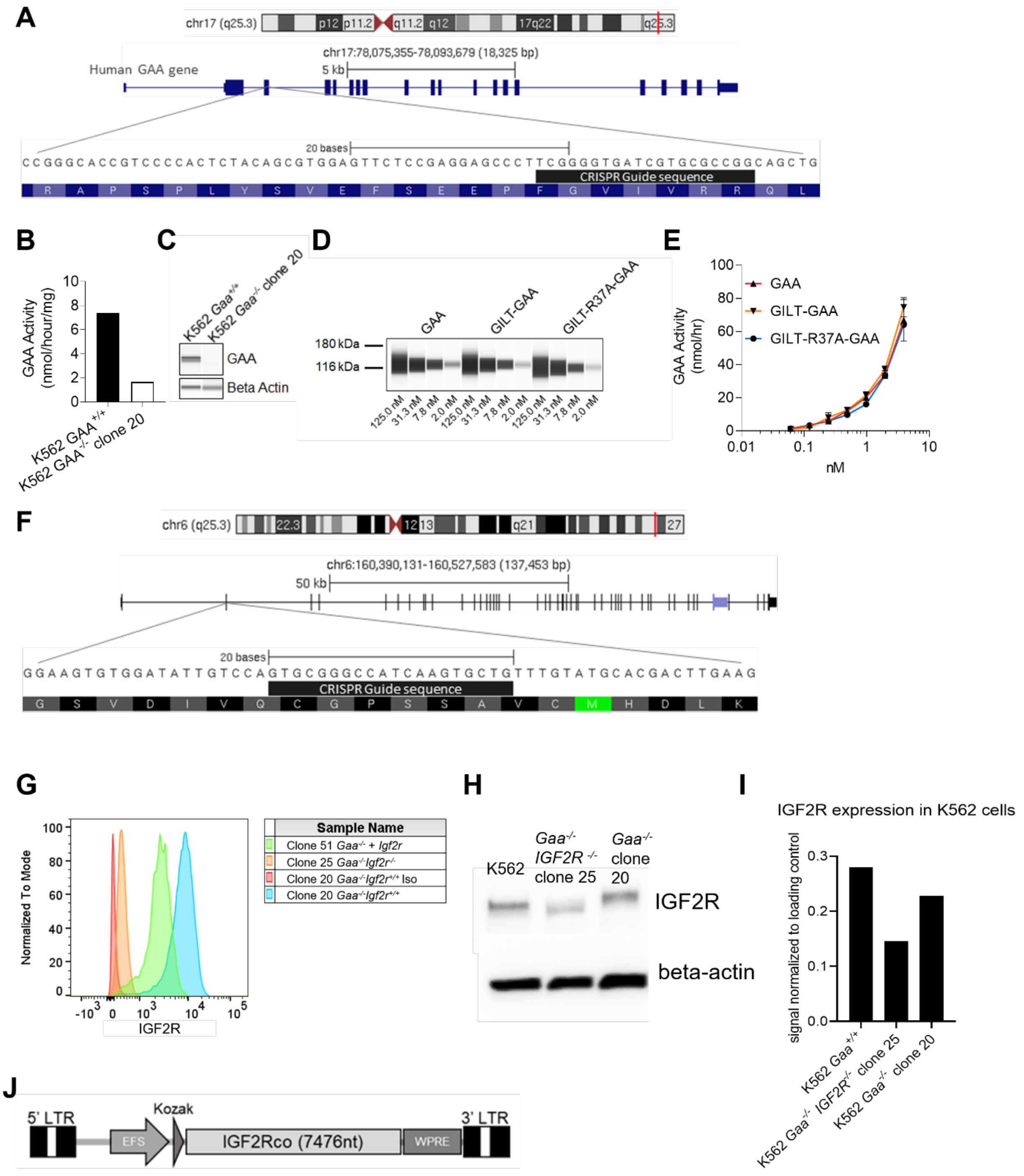


**Fig. S3. Characterization of gene-edited K562 GAA and IGF2R knockout cells.**

(**A**) Representation of the human *GAA* gene exon 3 specific CRISPR/Cas9 guide sequence aligned to the human reference genome hg19. (**B**) GAA activity of wildtype $GAA^{+/+}$ K562 cell lysates (ATCC) and $GAA^{-/-}$ K562 clone 20. (**C**) WB analysis of wildtype $GAA^{+/+}$ K562 cells and $GAA^{-/-}$ K562 clone 20 lysates using anti-GAA antibody (clone 1C12C11/F9). (**D**) WB of GAA, GILT-GAA and GILT-R37A-GAA protein preparations are presented in Fig. 1C and

D. (**E)** GAA activity of the three purified GAA, GILT-GAA and GILT-R37A-GAA protein preparations mentioned under Fig. S3E. (**F)** Representation of the human *IGF2R* gene exon 2 specific CRISPR/Cas9 guide sequence aligned to the human reference genome hg19. (**G**) Flow cytometry analysis of wild-type (WT) parental, *GAA*$^{-/-}$ clone 20, *GAA*$^{-/-}$ *IGF2R*$^{-/-}$ clone 25 and *GAA*$^{-/-}$ + *IGF2R* vector clone 51 K562 cells. (**H)** WB of WT, *GAA*$^{-/-}$ clone 20, *GAA*$^{-/-}$ *IGF2R*$^{-/-}$ clone 25 using an IGF2R specific monoclonal antibody. **(I)** Quantification of IGF2R protein via WB normalized to beta-actin loading control. (**J**) Representative schema of a lentiviral vector containing the codon optimized *IGF2R* transgene driven by the EFS promoter used to transduce *GAA*$^{-/-}$ *IGF2R*$^{-/-}$ clone 25 at MOI 10. The resulting *GAA*$^{-/-}$ + *IGF2R* add-back cell line was noted as clone 51.

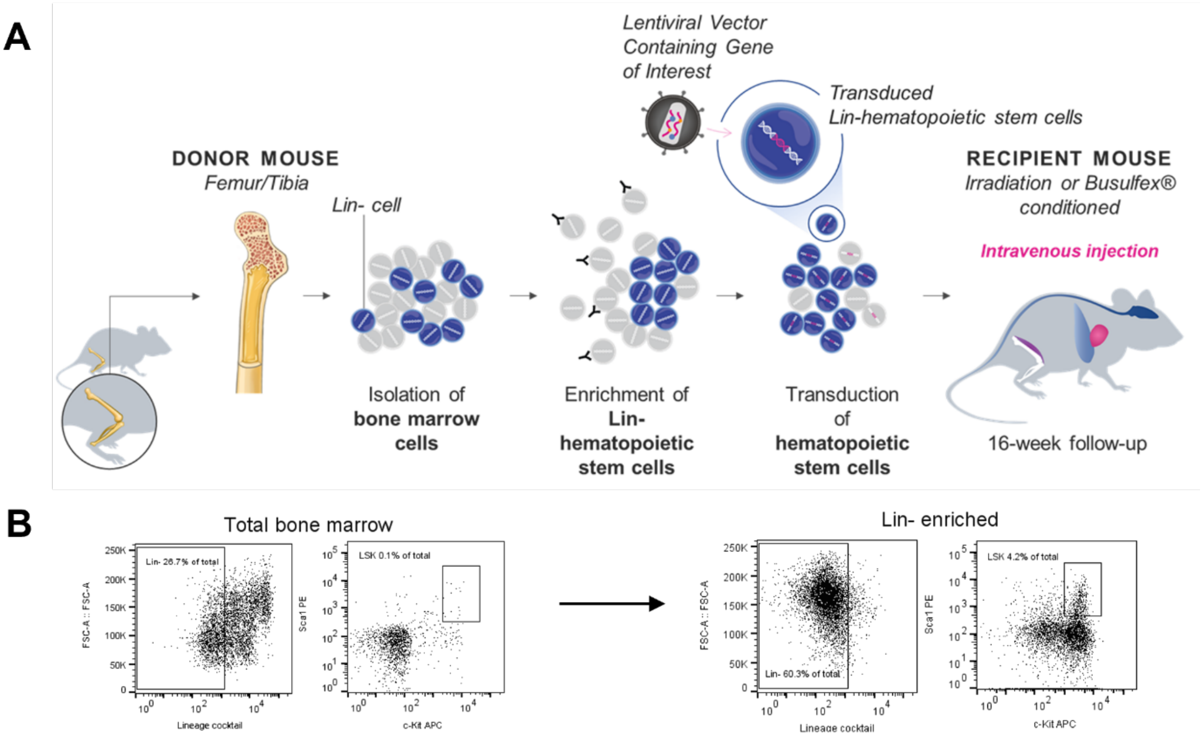


**Fig. S4. Illustration of experimental enrichment and transplant model.**

**(A)** Graphical depiction of the study design. Harvested donor bone marrow cells from femurs and tibias were Lin depleted using Robosep$^{TM}$ (StemCell Technologies), transduced with lentiviral vectors and infused into pre-conditioned recipient mice. (**B**) Representative dot plot of Lin- bone marrow cells enrichment. Frequencies of cell subpopulations are indicated as a percentage of total cells.

### Supplementary figure 5 part 1

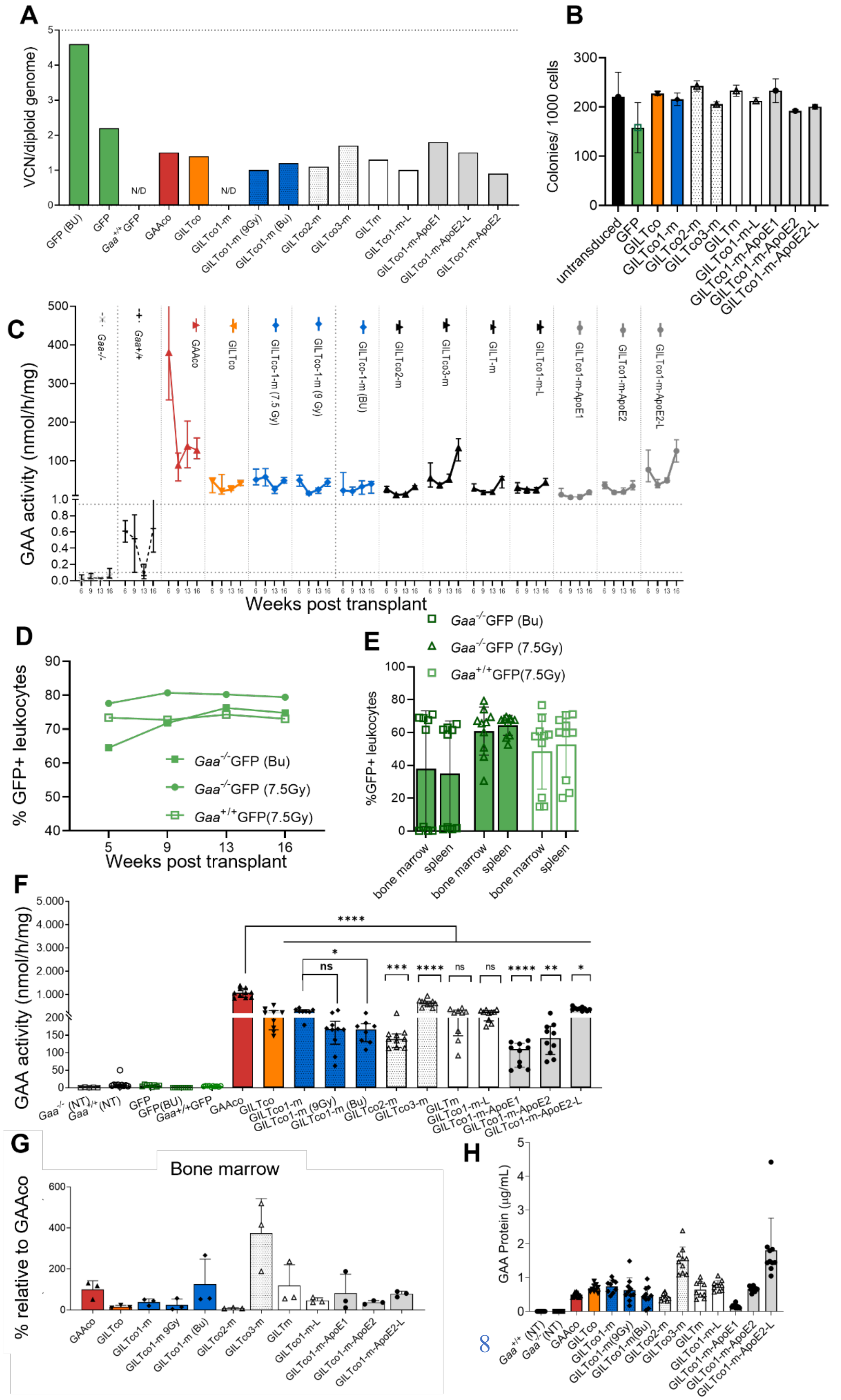

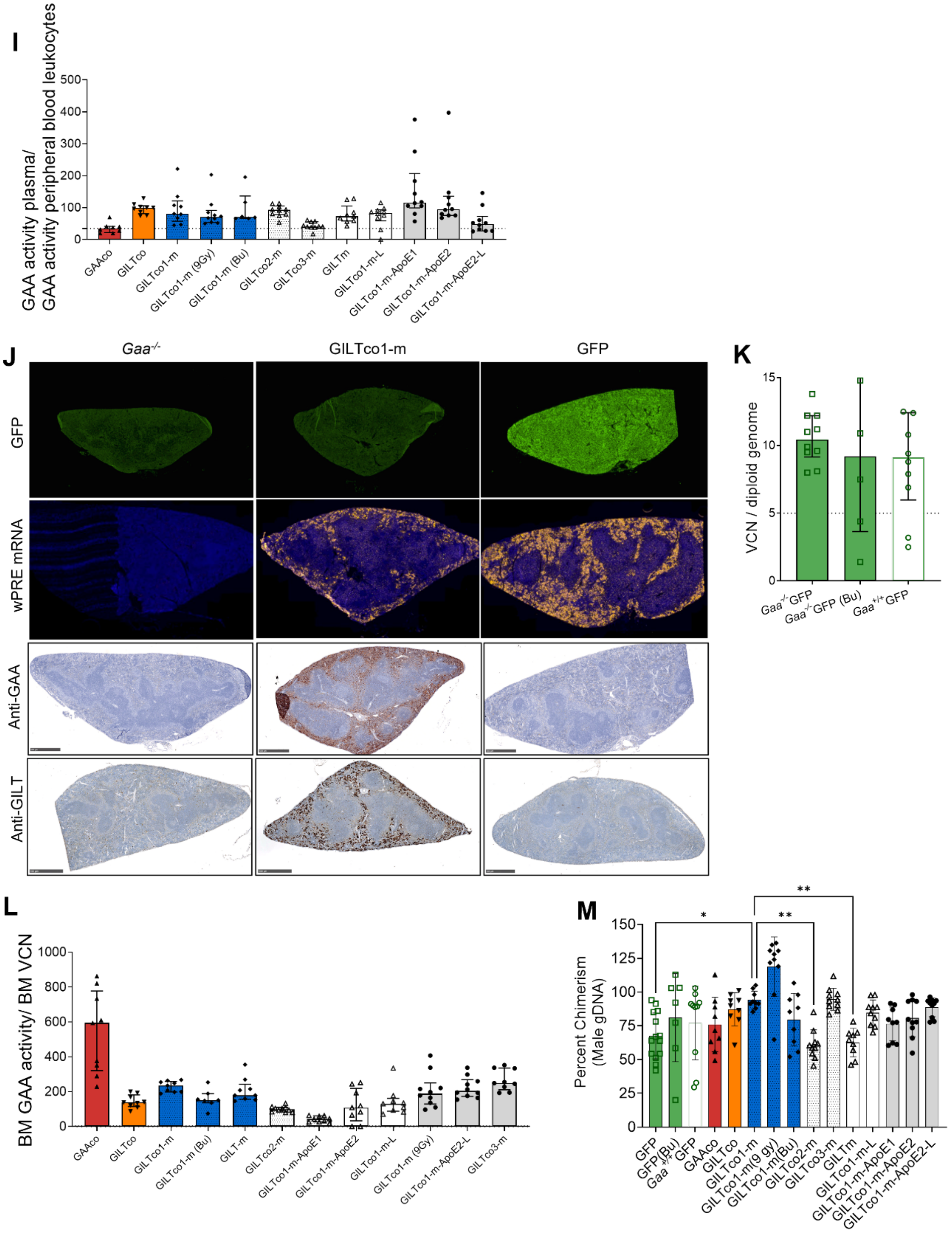
I
GAA activity plasma/
GAA activity peripheral blood leukocytes
GAAco
GILTco
GILTco1-m
GILTco1-m (9Gy)
GILTco1-m (Bu)
GILTco2-m
GILTco3-m
GILTm
GILTco1-m-L
GILTco1-m-ApoE1
GILTco1-m-ApoE2
GILTco1-m-ApoE2-L
J
Gaa-/-
GILTco1-m
GFP
wPRE mRNA
Anti-GAA
Anti-GILT
K
VCN / diploid genome
Gaa-/-GFP
Gaa-/-GFP (Bu)
Gaa+/+GFP
L
BM GAA activity/ BM VCN
GAAco
GILTco
GILTco1-m
GILTco1-m (Bu)
GILT-m
GILTco2-m
GILTco1-m-ApoE1
GILTco1-m-ApoE2
GILTco1-m-L
GILTco1-m (9Gy)
GILTco1-m-ApoE2-L
GILTco3-m
M
Percent Chimerism
(Male gDNA)
GFP
GFP(Bu)
Gaa+/+GFP
GAAco
GILTco
GILTco1-m
GILTco1-m(9 gy)
GILTco1-m(Bu)
GILTco2-m
GILTco3-m
GILTm
GILTco1-m-L
GILTco1-m-ApoE1
GILTco1-m-ApoE2
GILTco1-m-ApoE2-L

**Fig. S5. Characterization of genetically modified cultured donor cells and recipient hematopoietic compartment.**

Cultured transduced *Gaa*$^{-/-}$ Lin- donor bone marrow cells on day 7 were used to determine **(A)** VCN (**B)** and HSPC differentiation potentials in CFU assays (n=2). (**C**) GAA activity in PB leukocytes of *Gaa*$^{-/-}$ mice at weeks 6, 9, 13 and 16, group medians and interquartile ranges (n=10). (**D)** % GFP+ cells in PB CD45+ leukocytes, at weeks 5, 9, 13 and 16 post-transplant, group medians shown (n = 8-13). (**E**) % GFP+ CD45+ leukocytes in bone marrow and spleen of the GFP vector groups at 16 weeks post-transplant, individual values, means, and SD shown (n=9-10). (**F)** GAA activity in spleen at week 16 post-transplant, individual values, group medians, and interquartile ranges shown. Exact Wilcoxon Rank Sum p-values comparing arms with GAAco and with GILTco1-m (n=8-10). (**G)** Bone marrow cells GAA protein normalized to beta-actin loading control reported as % of GAAco group. **(H)** Final plasma GAA protein quantification per mL of mouse plasma (µg/mL). (**I**) Ratio of GAA activity in the plasma and PB leukocytes at 16 weeks post-transplant, medians and interquartile ranges shown. Dotted line is the average of the GAAco group used as a reference (n = 8-10). (**J**) GFP immunofluorescence (top), FISH *WPRE* (second row), GAA (third row) immunohistochemistry and GILT (bottom) transcripts in spleens. (**K)** VCN in bone marrow cell pellets measured at week 16 post-transplant**,** individual values, group medians and interquartile ranges (n = 5-10). **(L)** Bone marrow GAA activity normalized to bone marrow VCN (n=9-10) **(M)** Donor cells (male) chimerism via Y-chromosome PCR quantification, individual values, group means, and SD (n=7-10). Kruskal Wallis p-values GILTco1-m comparison to all treated groups, significant, where p-value *p< 0.05, **p<0.01, ***p<0.001 and ****p<0.0001.

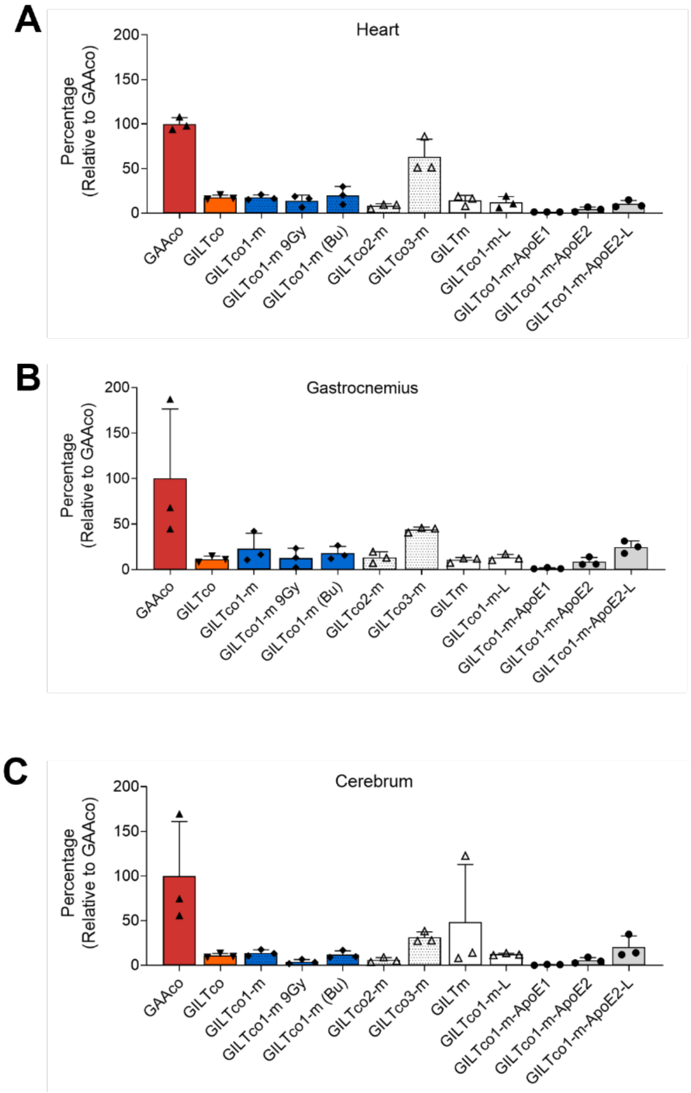


**Fig. S6. Relative GAA protein levels in heart, gastrocnemius and cerebrum in treatment groups.**

GAA protein expression levels in WB from **(A)** heart, **(B)** gastrocnemius and **(C)** cerebrum lysates normalized to GAPDH or beta-actin controls and reported as percentages of the respective GAAco group value.

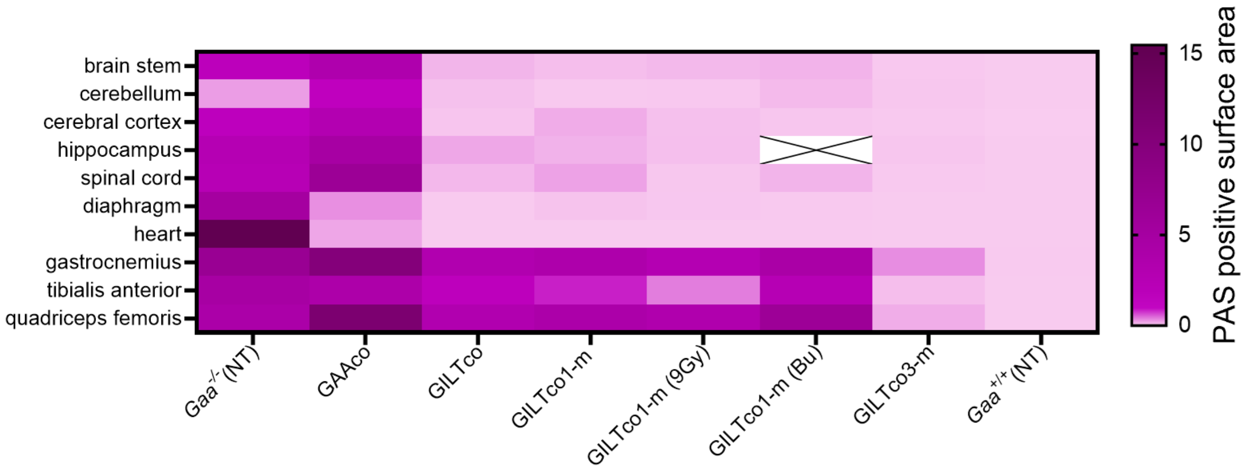


**Fig. S7. Quantification of glycogen via PAS staining in heart, skeletal muscles and CNS**.

Glycogen accumulation in tissues analyzed by PAS staining at 16 weeks post-transplant. Intensity (light to solid) of purple color depicts means of PAS+ surface area in representative tissue samples (n=3).

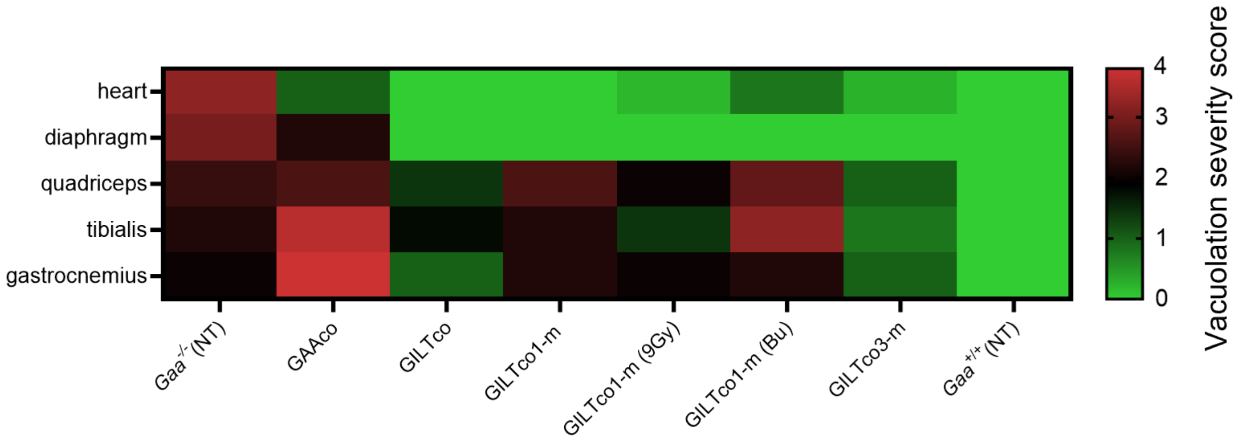


**Fig. S8. Quantification of myofiber vacuolation in heart and skeletal muscle.**

Assessment of selected groups for myofiber vacuolation in tissues stained with H&E. Severity scores assigned 0 to 4 from low to high, stained by H&E (n = 4-5).

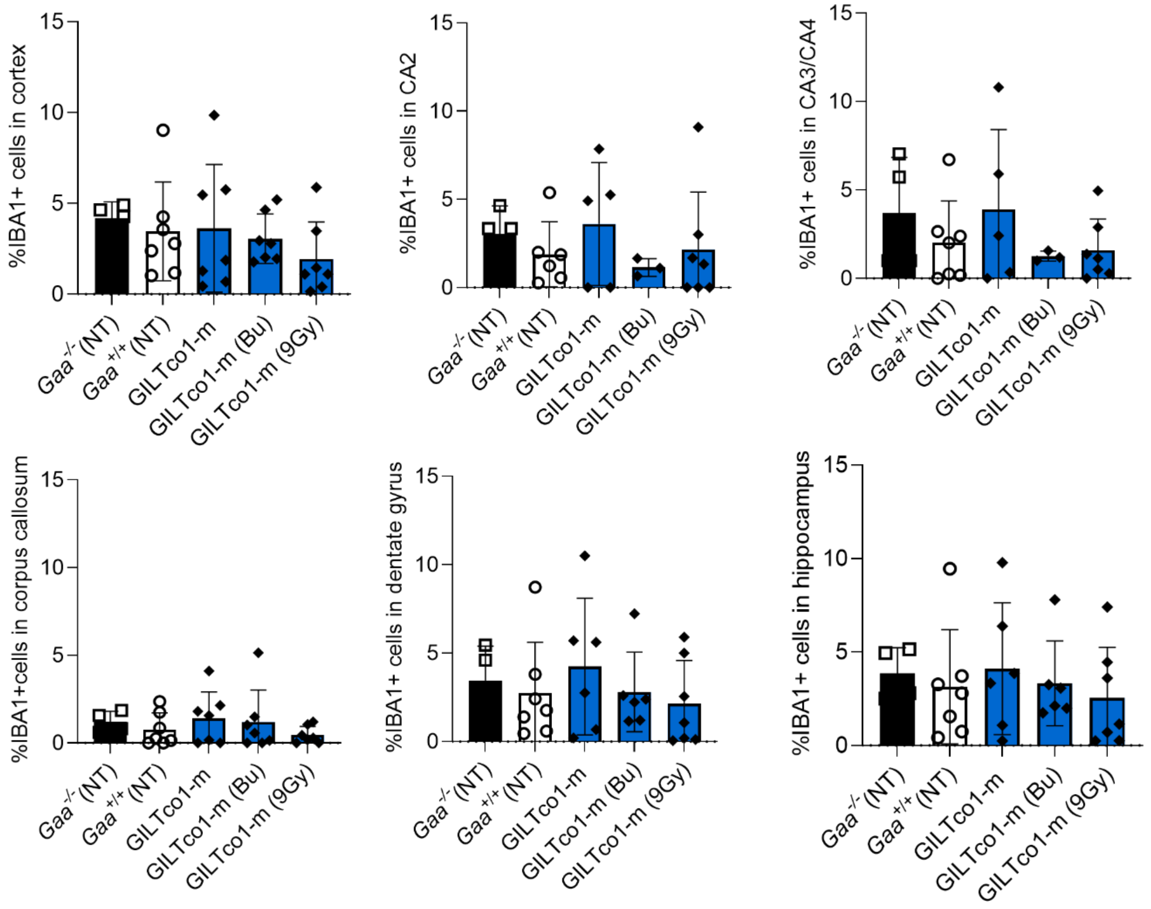


**Fig. S9. Quantification of total Iba+ cells in multiple brain regions.**

Brain tissue was stained for Iba1 and scanned slides were quantified by defining regions of interest where Iba1+ cells were counted as % of DAPI+ total cells per region. Group means with SD shown (n = 4-7).

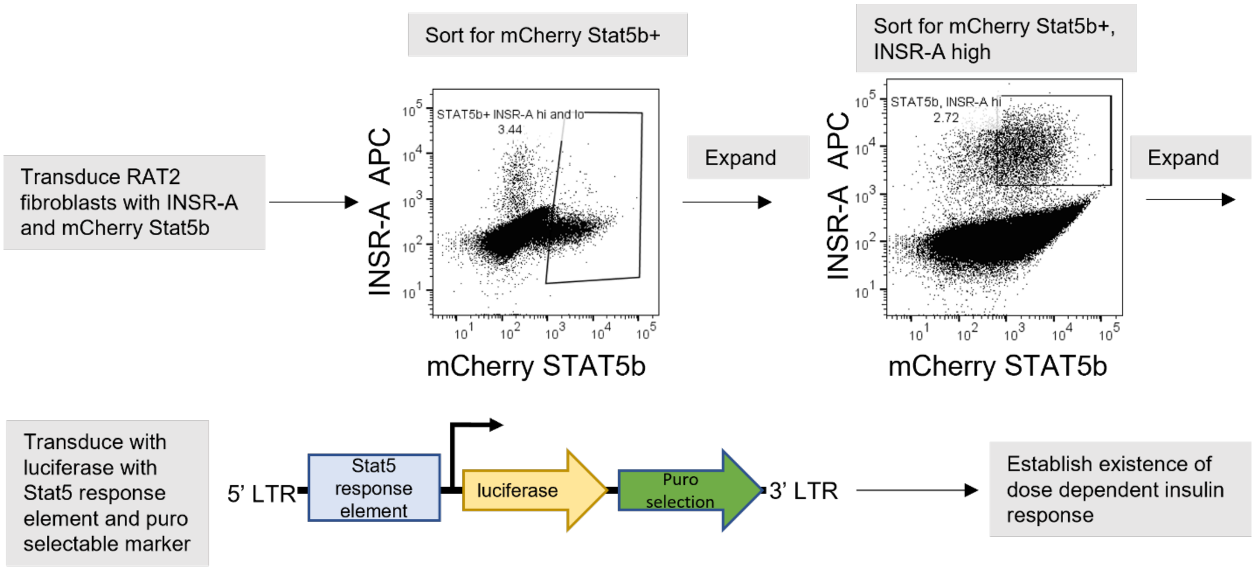


**Fig. S10. Diagram presenting the generation of insulin reporter cell line.**

RAT2 fibroblasts were transduced with a lentiviral vector expressing INSR-A and another expressing STAT5b, and sorted for INSR-A and mCherry positive cells. After expansion, the double positive INSR-A high/STAT5b high cells were collected, expanded, and transduced with a luciferase STAT5 reporter containing a puromycin selectable marker. Puromycin positive cells were selected and tested in a dose dependent luciferase response to insulin assay.

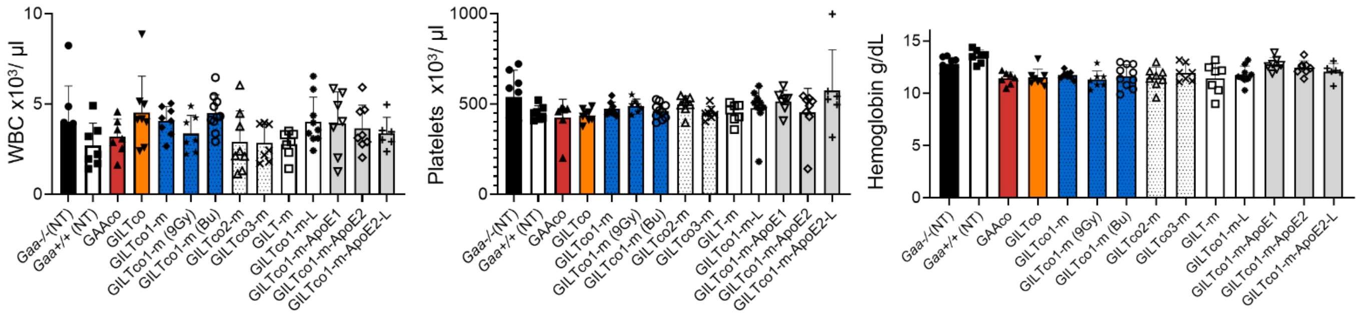


**Fig. S11. Complete blood count results in transplanted mice.**

Absolute counts of (**A**) white blood cells (**B**) platelets (**C**) hemoglobin at week 16 post-transplant. Individual values, group means, and SD shown (n = 6-9).

**Table S1 Study Design of the Vector Comparison Study.**

| N of Mice | Donor Phenotype | Recipient Phenotype | Abbreviated Transduced Cell Identity | Donor Cell Dose | Conditioning |
|---|---|---|---|---|---|
| 16 | N/A | *Gaa*$^{-/-}$ | *Gaa*$^{-/-}$ (NT) | N/A | N/A |
| 16 | N/A | *Gaa*$^{-/-}$ | *Gaa*$^{+/+}$ (NT) | N/A | N/A |
| 13 | *Gaa*$^{-/-}$ | *Gaa*$^{-/-}$ | GFP | 5 x $10^5$ | 7.5 Gy |
| 10 | *Gaa*$^{-/-}$ | *Gaa*$^{-/-}$ | GFP (Bu) | 5 x $10^5$ | Busulfex® |
| 10 | *Gaa*$^{+/+}$ | *Gaa*$^{+/+}$ | *Gaa*$^{+/+}$GFP | 5 x $10^5$ | 7.5 Gy |
| 10 | *Gaa*$^{-/-}$ | *Gaa*$^{-/-}$ | GAAco | 5 x $10^5$ | 7.5 Gy |
| 10 | *Gaa*$^{-/-}$ | *Gaa*$^{-/-}$ | GILTco | 5 x $10^5$ | 7.5 Gy |
| 10 | *Gaa*$^{-/-}$ | *Gaa*$^{-/-}$ | GILTco1-m | 5 x $10^5$ | 7.5 Gy |
| 10 | *Gaa*$^{-/-}$ | *Gaa*$^{-/-}$ | GILTco1-m (9Gy) | 5 x $10^5$ | 9.0 Gy |
| 10 | *Gaa*$^{-/-}$ | *Gaa*$^{-/-}$ | GILTco1-m (Bu) | 5 x $10^5$ | Busulfex® |
| 10 | *Gaa*$^{-/-}$ | *Gaa*$^{-/-}$ | GILTco2-m | 5 x $10^5$ | 7.5 Gy |
| 10 | *Gaa*$^{-/-}$ | *Gaa*$^{-/-}$ | GILTco3-m | 5 x $10^5$ | 7.5 Gy |
| 10 | *Gaa*$^{-/-}$ | *Gaa*$^{-/-}$ | GILTm | 5 x $10^5$ | 7.5 Gy |
| 10 | *Gaa*$^{-/-}$ | *Gaa*$^{-/-}$ | GILTco1-m-L | 5 x $10^5$ | 7.5 Gy |
| 10 | *Gaa*$^{-/-}$ | *Gaa*$^{-/-}$ | GILTco1-m-ApoE1 | 5 x $10^5$ | 7.5 Gy |
| 10 | *Gaa*$^{-/-}$ | *Gaa*$^{-/-}$ | GILTco1-m-ApoE2 | 5 x $10^5$ | 7.5 Gy |
| 10 | *Gaa*$^{-/-}$ | *Gaa*$^{-/-}$ | GILTco1-m-ApoE2-L | 5 x $10^5$ | 7.5 Gy |

**Table S2. PAS+ stain quantification in CNS and skeletal muscle.**

| | *Gaa*$^{-/-}$(NT) | *Gaa*$^{+/+}$(NT) | GAAco | GILTco | GILTco1-m | GILTco1-m (9Gy) | GILTco1-m (Bu) | GILTco3-m |
|---|---|---|---|---|---|---|---|---|
| **Brain Stem, N** | **3** | **3** | **3** | **3** | **3** | **3** | **3** | **3** |
| Percent PAS+, % | 1.6683 | 0.0011 | 3.6396 | 0.1558 | 0.0744 | 0.0816 | 0.7902 | 0.0140 |
| (SD) | (0.3424) | (0.0012) | (0.5917) | (0.0960) | (0.0788) | (0.0205) | (1.1699) | (0.0060) |
| **Cerebellum, N** | **3** | **3** | **3** | **3** | **3** | **3** | **3** | **2** |
| Percent PAS+, % | 0.2819 | 0.0016 | 1.2362 | 0.0756 | 0.0126 | 0.0174 | 0.2930 | 0.0223 |
| (SD) | (0.1137) | (0.0008) | (0.6042) | (0.0421) | (0.0119) | (0.0180) | (0.3941) | (0.0249) |
| **Cerebral Cortex, N** | **3** | **3** | **3** | **3** | **3** | **3** | **3** | **3** |
| Percent PAS+, % | 1.1863 | 0.0002 | 3.3613 | 0.0675 | 0.1259 | 0.0555 | 0.6911 | 0.0092 |
| (SD) | (0.4570) | (0.0002) | (1.1267) | (0.0603) | (0.1090) | (0.0094) | (1.1698) | (0.0044) |
| **Diaphragm, N** | **3** | **3** | **3** | **3** | **3** | **3** | **2** | **3** |
| Percent PAS+, % | 4.8662 | 0.0240 | 0.3120 | 0.0073 | 0.0309 | 0.0384 | 0.0106 | 0.0072 |
| (SD) | (2.7952) | (0.0404) | (0.1690) | (0.0045) | (0.0248) | (0.0358) | (0.0040) | (0.0097) |
| **Gastrocnemius, N** | **3** | **3** | **3** | **3** | **3** | **3** | **2** | **3** |
| Percent PAS+, % | 6.8983 | 0.0111 | 10.0793 | 2.5953 | 3.8556 | 4.2678 | 4.3241 | 0.3160 |
| (SD) | (4.5706) | (0.0090) | (1.1000) | (1.6335) | (1.3688) | (4.1667) | (3.1180) | (0.0928) |
| **Heart, N** | **3** | **3** | **3** | **3** | **3** | **3** | **3** | **3** |
| Percent PAS+, % | 14.9640 | 0.0016 | 0.3090 | 0.0025 | 0.0037 | 0.0026 | 1.7620 | 0.0005 |
| (SD) | (9.1151) | (0.0021) | (0.2585) | (0.0018) | (0.0034) | (0.0006) | (3.0408) | (0.0004) |
| **Hippocampus, N** | **1** | **2** | **2** | **3** | **3** | **3** | **0** | **2** |
| Percent PAS+, % | 2.5743 | 0.0009 | 4.7405 | 0.1680 | 0.1584 | 0.0675 | N/A | 0.0295 |
| (SD) | N/A | (0.0012) | (2.6534) | (0.0829) | (0.1703) | (0.0327) | N/A | (0.0102) |
| **Quadriceps Femoris, N** | **3** | **3** | **3** | **3** | **3** | **3** | **3** | **3** |
| Percent PAS+, % | 8.8374 | 0.0725 | 10.9931 | 2.1886 | 3.7669 | 2.4127 | 7.2275 | 0.1554 |
| (SD) | (10.3491) | (0.1227) | (0.8839) | (1.8304) | (0.6523) | (1.9177) | (5.5191) | (0.0356) |
| **Spinal Cord, N** | **3** | **3** | **3** | **3** | **3** | **3** | **3** | **3** |
| Percent PAS+, % | 2.2401 | 0.0028 | 5.8312 | 0.1205 | 0.2225 | 0.0256 | 1.8864 | 0.0142 |
| (SD) | (0.4483) | (0.0017) | (0.9861) | (0.0706) | (0.0742) | (0.0131) | (3.1222) | (0.0041) |
| **Tibialis Anterior, N** | **3** | **3** | **3** | **3** | **3** | **3** | **3** | **3** |
| Percent PAS+, % | 5.5348 | 0.0086 | 3.5480 | 0.9759 | 1.2340 | 0.4417 | 6.9388 | 0.0570 |
| (SD) | (4.4465) | (0.0094) | (3.1285) | (0.7629) | (0.8260) | (0.4188) | (8.7839) | (0.0368) |

**Table S3. Vacuolation scores in heart and skeletal muscle.**

| | *Gaa*$^{-/-}$(NT) | *Gaa*$^{+/+}$(NT) | GAAco | GILTco | GILTco1-m | GILTco1-m (9Gy) | GILTco1-m (Bu) | GILTco3-m |
|---|---|---|---|---|---|---|---|---|
| **Heart (# examined)** | **5** | **5** | **5** | **5** | **5** | **5** | **5** | **5** |
| **Vacuolation, myofiber** | **5** | **0** | **5** | **0** | **0** | **0** | **3** | **1** |
| Minimal | 0 | 0 | 5 | 0 | 0 | 0 | 2 | 1 |
| Mild | 0 | 0 | 0 | 0 | 0 | 0 | 1 | 0 |
| Moderate | 4 | 0 | 0 | 0 | 0 | 0 | 0 | 0 |
| Marked | 1 | 0 | 0 | 0 | 0 | 0 | 0 | 0 |
| **Vacuolation, vascular** | **5** | **0** | **3** | **0** | **0** | **0** | **1** | **0** |
| Minimal | 2 | 0 | 3 | 0 | 0 | 0 | 1 | 0 |
| Mild | 3 | 0 | 0 | 0 | 0 | 0 | 0 | 0 |
| **Quadriceps femoris (# examined)** | **5** | **5** | **5** | **5** | **5** | **5** | **5** | **5** |
| **Vacuolation, myofiber** | **5** | **0** | **5** | **5** | **5** | **5** | **5** | **5** |
| Minimal | 0 | 0 | 0 | 3 | 0 | 0 | 0 | 5 |
| Mild | 3 | 0 | 0 | 2 | 2 | 5 | 1 | 0 |
| Moderate | 2 | 0 | 3 | 0 | 3 | 0 | 4 | 0 |
| Marked | 0 | 0 | 2 | 0 | 0 | 0 | 0 | 0 |
| **Vacuolation, vascular** | **1** | **0** | **2** | **0** | **0** | **0** | **0** | **0** |
| Minimal | 1 | 0 | 2 | 0 | 0 | 0 | 0 | 0 |
| **Diaphragm (# examined)** | **5** | **5** | **5** | **5** | **5** | **5** | **4** | **5** |
| **Vacuolation, myofiber** | **5** | **0** | **5** | **0** | **0** | **0** | **0** | **0** |
| Minimal | 0 | 0 | 0 | 0 | 0 | 0 | 0 | 0 |
| Mild | 0 | 0 | 4 | 0 | 0 | 0 | 0 | 0 |
| Moderate | 5 | 0 | 1 | 0 | 0 | 0 | 0 | 0 |
| **Vacuolation, vascular** | **4** | **0** | **1** | **0** | **0** | **0** | **0** | **0** |
| Minimal | 0 | 0 | 0 | 0 | 0 | 0 | 0 | 0 |
| Mild | 2 | 0 | 1 | 0 | 0 | 0 | 0 | 0 |
| Moderate | 2 | 0 | 0 | 0 | 0 | 0 | 0 | 0 |
| **Gastrocnemius (# examined)** | **5** | **5** | **5** | **5** | **5** | **5** | **5** | **5** |
| **Vacuolation, myofiber** | **5** | **0** | **5** | **5** | **5** | **5** | **4** | **5** |
| Minimal | 0 | 0 | 0 | 0 | 0 | 0 | 0 | 5 |
| Mild | 5 | 0 | 0 | 5 | 4 | 5 | 1 | 0 |
| Moderate | 0 | 0 | 1 | 0 | 1 | 0 | 3 | 0 |
| Marked | 0 | 0 | 4 | 0 | 0 | 0 | 0 | 0 |
| **Vacuolation, vascular** | **4** | **0** | **2** | **0** | **0** | **0** | **0** | **0** |
| Minimal | 0 | 0 | 2 | 0 | 0 | 0 | 0 | 0 |
| Mild | 4 | 0 | 0 | 0 | 0 | 0 | 0 | 0 |
| **Tibialis anterior (# examined)** | **5** | **5** | **5** | **5** | **5** | **5** | **5** | **5** |
| **Vacuolation, myofiber** | **5** | **0** | **5** | **5** | **5** | **5** | **5** | **4** |
| Minimal | 0 | 0 | 0 | 1 | 0 | 3 | 0 | 4 |
| Mild | 4 | 0 | 0 | 4 | 4 | 2 | 0 | 0 |
| Moderate | 1 | 0 | 2 | 0 | 1 | 0 | 4 | 0 |
| Marked | 0 | 0 | 3 | 0 | 0 | 0 | 1 | 0 |
| **Vacuolation, vascular** | **5** | **0** | **2** | **0** | **0** | **0** | **1** | **0** |
| Minimal | 0 | 0 | 0 | 0 | 0 | 0 | 1 | 0 |
| Mild | 5 | 0 | 0 | 0 | 0 | 0 | 0 | 0 |
| Moderate | 0 | 0 | 0 | 0 | 0 | 0 | 0 | 0 |
| Marked | 0 | 0 | 2 | 0 | 0 | 0 | 0 | 0 |

**Table S4 Vacuolation and degeneration scores in CNS**

| | *Gaa*$^{-/-}$(NT) | *Gaa*$^{+/+}$(NT) | GAAco | GILTco | GILTco1-m | GILTco1-m (Bu) | GILTco1-m (9Gy) | GILTco3-m |
|---|---|---|---|---|---|---|---|---|
| **Brain (# examined)** | **5** | **5** | **5** | **5** | **5** | **5** | **5** | **5** |
| **Vacuolation, neuronal/axonal** | **5** | **5** | **5** | **5** | **5** | **5** | **5** | **3** |
| Minimal | 0 | 0 | 0 | 5 | 3 | 2 | 4 | 3 |
| Mild | 0 | 0 | 0 | 0 | 2 | 1 | 1 | 0 |
| Moderate | 1 | 0 | 0 | 0 | 0 | 2 | 0 | 0 |
| Marked | 4 | 0 | 5 | 0 | 0 | 0 | 0 | 0 |
| **Degeneration/necrosis neuronal/axonal** | **5** | **0** | **1** | **4** | **3** | **4** | **4** | **4** |
| Minimal | 3 | 0 | 0 | 1 | 2 | 3 | 2 | 2 |
| Mild | 2 | 0 | 1 | 3 | 1 | 1 | 2 | 2 |
| **Vacuolation, vascular** | **5** | **0** | **5** | **1** | **4** | **4** | **0** | **0** |
| Minimal | 1 | 0 | 1 | 1 | 3 | 0 | 0 | 0 |
| Mild | 3 | 0 | 4 | 0 | 1 | 4 | 0 | 0 |
| Moderate | 1 | 0 | 0 | 0 | 0 | 0 | 0 | 0 |
| **Vacuolation, meninges** | **4** | **0** | **0** | **0** | **0** | **1** | **0** | **0** |
| Minimal | 3 | 0 | 0 | 0 | 0 | 1 | 0 | 0 |
| Mild | 1 | 0 | 0 | 0 | 0 | 0 | 0 | 0 |
| **Vacuolation, choriod plexus** | **2** | **0** | **2** | **3** | **1** | **4** | **0** | **0** |
| Minimal | **2** | **0** | **2** | **3** | **1** | **4** | **0** | **0** |
| **Spinal Cord (# examined)** | **5** | **5** | **5** | **5** | **5** | **5** | **5** | **5** |
| **Vacuolation, neuronal/axonal** | **5** | **0** | **5** | **1** | **5** | **5** | **0** | **1** |
| Minimal | 0 | 0 | 0 | 1 | 5 | 4 | 0 | 1 |
| Mild | 0 | 0 | 0 | 0 | 0 | 0 | 0 | 0 |
| Moderate | 1 | 0 | 0 | 0 | 0 | 0 | 0 | 0 |
| Marked | **4** | **0** | **5** | **0** | **0** | **0** | **0** | **0** |
| Severe | 0 | 0 | 0 | 0 | 0 | 1 | 0 | 0 |
| **Degeneration/necrosis neuronal/axonal** | **3** | **0** | **1** | **2** | **2** | **2** | **0** | **1** |
| Minimal | 1 | 0 | 1 | 2 | 2 | 1 | 0 | 1 |
| Mild | 2 | 0 | 0 | 0 | 0 | 1 | 0 | 0 |
| **Vacuolation, vascular** | **2** | **0** | **1** | **0** | **0** | **0** | **0** | **0** |
| Mild | 2 | 0 | 1 | 0 | 0 | 0 | 0 | 0 |

**Table S5. Flow cytometry immunophenotyping of the PB leukocytes at week 16 post-transplant.**

| PB leukocytes 16 weeks | | | | | | | |
|---|---|---|---|---|---|---|---|
| Group name | | % total leukocytes | % GR1+ Mac1+ | % B220+ | %CD3+ | %CD4+ | %CD8+ |
| *Gaa*$^{-/-}$ | n | 6 | 6 | 6 | 6 | 6 | 6 |
| | Mean | 91.61 | 29.04 | 33.98 | 24.39 | 16.46 | 7.21 |
| | SD | 3.01 | 8.68 | 6.85 | 5.57 | 4.38 | 1.70 |
| *Gaa*$^{+/+}$ | n | 9 | 9 | 9 | 9 | 9 | 9 |
| | Mean | 91.45 | 31.20 | 32.14 | 30.15 | 20.00 | 9.30 |
| | SD | 3.85 | 9.26 | 10.09 | 8.07 | 6.27 | 2.09 |
| GFP | n | 7 | 7 | 7 | 7 | 7 | 7 |
| | Mean | 96.10 | 15.43 | 34.67 | 40.75 | 29.41 | 10.57 |
| | SD | 1.40 | 4.14 | 2.81 | 2.94 | 2.53 | 1.69 |
| GFP (Bu) | n | 5 | 5 | 5 | 5 | 5 | 5 |
| | Mean | 94.87 | 23.17 | 30.97 | 38.02 | 28.88 | 8.51 |
| | SD | 2.35 | 9.25 | 5.04 | 6.66 | 5.42 | 1.63 |
| GFP *Gaa*$^{+/+}$ | n | 6 | 6 | 6 | 6 | 6 | 6 |
| | Mean | 94.42 | 12.87 | 35.14 | 44.03 | 32.17 | 10.89 |
| | SD | 1.84 | 3.83 | 4.96 | 5.77 | 4.91 | 1.74 |
| GAAco | n | 6 | 6 | 6 | 6 | 6 | 6 |
| | Mean | 92.97 | 36.55 | 28.45 | 28.57 | 21.25 | 6.83 |
| | SD | 1.98 | 11.21 | 8.29 | 3.82 | 2.52 | 1.81 |
| GILTco | n | 7 | 7 | 7 | 7 | 7 | 7 |
| | Mean | 93.08 | 33.22 | 35.50 | 25.13 | 17.36 | 7.23 |
| | SD | 1.08 | 11.05 | 10.17 | 6.88 | 4.90 | 2.05 |
| GILTco1-m | n | 9 | 9 | 9 | 9 | 9 | 9 |
| | Mean | 94.26 | 32.07 | 28.62 | 32.89 | 23.04 | 9.19 |
| | SD | 2.01 | 10.75 | 9.87 | 3.63 | 3.68 | 1.00 |
| GILTco1-m (9Gy) | n | 6 | 6 | 6 | 6 | 6 | 6 |
| | Mean | 92.20 | 25.94 | 35.77 | 30.61 | 23.29 | 6.81 |
| | SD | 0.89 | 4.19 | 4.85 | 2.58 | 2.54 | 0.29 |
| GILTco1-m (Bu) | n | 10 | 10 | 10 | 10 | 10 | 10 |
| | Mean | 93.26 | 43.96 | 23.24 | 27.83 | 19.47 | 7.77 |
| | SD | 1.88 | 7.94 | 8.00 | 4.86 | 3.11 | 2.01 |
| GILTco2-m | n | 8 | 8 | 8 | 8 | 8 | 8 |
| | Mean | 91.32 | 25.68 | 32.91 | 31.59 | 23.77 | 7.25 |
| | SD | 2.11 | 6.53 | 8.80 | 4.90 | 4.38 | 1.28 |
| GILTco3-m | n | 7 | 7 | 7 | 7 | 7 | 7 |
| | Mean | 89.47 | 29.87 | 29.73 | 31.13 | 23.50 | 7.01 |
| | SD | 2.92 | 8.73 | 4.41 | 5.34 | 4.14 | 1.28 |
| GILT-m | n | 6 | 6 | 6 | 6 | 6 | 6 |
| | Mean | 88.97 | 27.40 | 32.64 | 30.14 | 23.50 | 6.07 |
| | SD | 2.96 | 7.81 | 8.40 | 5.59 | 4.81 | 0.98 |
| GILTco1-m-L | n | 8 | 8 | 8 | 8 | 8 | 8 |
| | Mean | 91.63 | 23.73 | 35.86 | 31.12 | 23.61 | 6.98 |
| | SD | 2.54 | 3.98 | 3.09 | 1.60 | 1.61 | 0.70 |
| GILTco1-m-ApoE1 | n | 7 | 7 | 7 | 7 | 7 | 7 |
| | Mean | 92.97 | 25.17 | 39.41 | 28.80 | 20.88 | 7.36 |
| | SD | 2.31 | 5.27 | 6.94 | 4.82 | 3.73 | 1.14 |
| GILTco1-m-ApoE2 | n | 5 | 5 | 5 | 5 | 5 | 5 |
| | Mean | 88.62 | 22.39 | 32.60 | 35.80 | 27.23 | 8.03 |
| | SD | 3.90 | 1.70 | 3.19 | 1.59 | 1.89 | 0.63 |
| GILTco1-m-ApoE2-L | n | 3 | 3 | 3 | 3 | 3 | 3 |
| | Mean | 83.85 | 26.37 | 29.91 | 35.21 | 26.64 | 7.94 |
| | SD | 7.21 | 6.13 | 4.97 | 2.17 | 2.77 | 0.65 |

**Table S6. Flow cytometry immunophenotyping of the bone marrow at week 16 post-transplant.**

| Group name | | %total leukocytes | % GR1+ Mac1+ | % B220+ | % CD3+ | % CD4+ | % CD8+ |
|---|---|---|---|---|---|---|---|
| ***Gaa***$^{-/-}$ | n | 10 | 10 | 10 | 10 | 10 | 10 |
| | Mean | 97.70 | 58.99 | 22.86 | 3.44 | 1.16 | 1.59 |
| | SD | 0.61 | 5.67 | 4.56 | 1.45 | 0.44 | 0.87 |
| ***Gaa***$^{+/+}$ | n | 10 | 10 | 10 | 10 | 10 | 10 |
| | Mean | 97.37 | 48.80 | 28.11 | 6.92 | 2.82 | 3.10 |
| | SD | 2.29 | 5.24 | 5.02 | 1.73 | 0.82 | 1.05 |
| **GFP** | n | 10 | 10 | 10 | 10 | 10 | 10 |
| | Mean | 97.82 | 33.20 | 41.95 | 8.83 | 3.38 | 4.34 |
| | SD | 0.38 | 7.46 | 6.59 | 2.17 | 1.05 | 1.16 |
| **GFP (Bu)** | n | 5 | 5 | 5 | 5 | 5 | 5 |
| | Mean | 97.70 | 38.13 | 38.80 | 7.91 | 3.28 | 3.62 |
| | SD | 0.59 | 6.32 | 5.58 | 1.23 | 0.58 | 0.67 |
| **GFP *Gaa***$^{+/+}$ | n | 10 | 10 | 10 | 10 | 10 | 10 |
| | Mean | 98.53 | 37.33 | 35.76 | 10.56 | 4.72 | 4.47 |
| | SD | 0.31 | 13.78 | 9.70 | 4.83 | 2.16 | 2.28 |
| **GAAco** | n | 10 | 10 | 10 | 10 | 10 | 10 |
| | Mean | 97.90 | 57.21 | 23.89 | 4.35 | 1.58 | 1.89 |
| | SD | 0.46 | 2.37 | 2.12 | 0.79 | 0.44 | 0.42 |
| **GILTco** | n | 9 | 9 | 9 | 9 | 9 | 9 |
| | Mean | 97.53 | 54.59 | 23.48 | 5.71 | 1.74 | 2.10 |
| | SD | 0.76 | 5.74 | 4.24 | 1.54 | 0.61 | 0.58 |
| **GILTco1-m** | n | 9 | 9 | 9 | 9 | 9 | 9 |
| | Mean | 97.33 | 55.94 | 23.94 | 4.39 | 1.60 | 1.86 |
| | SD | 0.51 | 2.31 | 2.28 | 0.88 | 0.43 | 0.43 |
| **GILTco1-m (9Gy)** | n | 10 | 10 | 10 | 10 | 10 | 10 |
| | Mean | 97.62 | 53.39 | 26.92 | 5.16 | 2.02 | 2.59 |
| | SD | 0.44 | 6.41 | 5.63 | 1.24 | 0.50 | 0.72 |
| **GILTco1-m (Bu)** | n | 10 | 10 | 10 | 10 | 10 | 10 |
| | Mean | 97.52 | 54.41 | 24.53 | 4.87 | 1.70 | 2.03 |
| | SD | 0.42 | 4.89 | 4.32 | 1.56 | 0.31 | 0.55 |
| **GILTco2-m** | n | 10 | 10 | 10 | 10 | 10 | 10 |
| | Mean | 97.92 | 53.35 | 27.75 | 4.52 | 1.72 | 2.26 |
| | SD | 0.39 | 4.34 | 4.10 | 1.13 | 0.48 | 0.60 |
| **GILTco3-m** | n | 10 | 10 | 10 | 10 | 10 | 10 |
| | Mean | 97.15 | 54.84 | 24.80 | 5.09 | 1.90 | 2.62 |
| | SD | 2.21 | 5.41 | 4.09 | 1.23 | 0.52 | 0.69 |
| **GILT-m** | n | 9 | 9 | 9 | 9 | 9 | 9 |
| | Mean | 97.94 | 52.91 | 27.11 | 5.43 | 2.10 | 2.71 |
| | SD | 0.26 | 2.39 | 2.69 | 0.95 | 0.39 | 0.56 |
| **GILTco1-m-L** | n | 10 | 10 | 10 | 10 | 10 | 10 |

| | | | | | | | |
|---|---|---|---|---|---|---|---|
| | Mean | 97.77 | 48.52 | 29.72 | 4.68 | 1.75 | 2.40 |
| | SD | 0.49 | 3.35 | 3.32 | 0.83 | 0.45 | 0.39 |
| **GILTco1-m-ApoE1** | n | 10 | 10 | 10 | 10 | 10 | 10 |
| | Mean | 98.04 | 52.80 | 28.36 | 4.70 | 1.91 | 2.28 |
| | SD | 0.35 | 4.94 | 4.59 | 0.61 | 0.31 | 0.32 |
| **GILTco1-m-ApoE2** | n | 10 | 10 | 10 | 10 | 10 | 10 |
| | Mean | 97.61 | 49.89 | 28.16 | 5.56 | 2.20 | 2.80 |
| | SD | 0.28 | 1.98 | 1.43 | 1.16 | 0.56 | 0.59 |
| **GILTco1-m-ApoE2-L** | n | 10 | 10 | 10 | 10 | 10 | 10 |
| | Mean | 97.66 | 51.64 | 26.90 | 5.78 | 2.23 | 2.94 |
| | SD | 0.37 | 4.57 | 3.54 | 1.33 | 0.52 | 0.76 |

**Table S7. Flow cytometry immunophenotyping of the spleen at week 16 post-transplant.**

| Group name | | % total leukocytes | % GR1+ Mac1+ | % B220+ | %CD3+ | %CD4+ | %CD8+ |
|---|---|---|---|---|---|---|---|
| **Gaa-/-** | n | 10 | 10 | 10 | 10 | 10 | 10 |
| | Mean | 96.13 | 5.96 | 55.63 | 26.35 | 13.60 | 10.73 |
| | SD | 3.16 | 3.56 | 4.63 | 5.53 | 3.41 | 2.47 |
| **Gaa+/+** | n | 10 | 10 | 10 | 10 | 10 | 10 |
| | Mean | 96.83 | 4.34 | 49.95 | 37.75 | 19.66 | 16.29 |
| | SD | 1.10 | 1.31 | 4.77 | 3.80 | 2.14 | 1.84 |
| **GFP** | n | 10 | 10 | 10 | 10 | 10 | 10 |
| | Mean | 98.58 | 2.80 | 57.80 | 31.40 | 18.44 | 10.87 |
| | SD | 0.45 | 0.63 | 3.37 | 3.63 | 2.39 | 1.19 |
| **GFP (Bu)** | n | 5 | 5 | 5 | 5 | 5 | 5 |
| | Mean | 98.85 | 2.73 | 56.92 | 32.95 | 19.64 | 11.32 |
| | SD | 0.24 | 0.85 | 3.03 | 3.52 | 2.09 | 1.59 |
| **GFP Gaa+/+** | n | 10 | 10 | 10 | 10 | 10 | 10 |
| | Mean | 98.83 | 1.98 | 58.12 | 33.09 | 20.00 | 11.27 |
| | SD | 0.68 | 1.52 | 4.62 | 3.31 | 2.92 | 1.13 |
| **GAAco** | n | 10 | 10 | 10 | 10 | 10 | 10 |
| | Mean | 97.29 | 6.31 | 49.55 | 34.51 | 22.07 | 11.02 |
| | SD | 1.16 | 2.32 | 3.93 | 4.69 | 3.31 | 1.43 |
| **GILTco** | n | | | | | 9 | 9 |
| | Mean | 97.66 | 5.58 | 56.10 | 30.47 | 18.13 | 10.94 |
| | SD | 0.77 | 2.53 | 5.67 | 6.04 | 4.41 | 1.92 |
| **GILTco1-m** | n | 9 | 9 | 9 | 9 | 9 | 9 |
| | Mean | 96.66 | 7.83 | 47.74 | 34.39 | 21.36 | 11.33 |
| | SD | 1.11 | 2.32 | 3.67 | 3.87 | 3.00 | 1.22 |
| **GILTco1-m (9Gy)** | n | 10 | 10 | 10 | 10 | 10 | 10 |
| | Mean | 97.48 | 5.19 | 55.50 | 31.03 | 19.56 | 10.02 |
| | SD | 0.94 | 1.16 | 2.06 | 2.03 | 1.64 | 0.95 |
| **GILTco1-m (Bu)** | n | | | | | 10 | 10 |

| | | | | | | | |
|---|---|---|---|---|---|---|---|
| | Mean | 98.40 | 4.70 | 54.65 | 32.35 | 19.55 | 11.45 |
| | SD | 0.32 | 1.20 | 3.11 | 3.34 | 2.33 | 1.24 |
| **GILTco2-m** | n | 10 | 10 | 10 | 10 | 10 | 10 |
| | Mean | 97.69 | 5.11 | 56.59 | 29.01 | 18.01 | 9.72 |
| | SD | 0.69 | 1.59 | 3.76 | 4.13 | 2.91 | 1.37 |
| **GILTco3-m** | n | 10 | 10 | 10 | 10 | 10 | 10 |
| | Mean | 96.95 | 5.13 | 54.31 | 30.79 | 19.25 | 10.26 |
| | SD | 0.73 | 2.00 | 2.81 | 3.18 | 2.36 | 1.56 |
| **GILT-m** | n | 9 | 9 | 9 | 9 | 9 | 9 |
| | Mean | 97.02 | 4.05 | 4.82 | 54.21 | 19.55 | 10.06 |
| | SD | 0.58 | 0.81 | 1.45 | 2.73 | 3.45 | 1.35 |
| **GILTco1-m-L** | n | 10 | 10 | 10 | 10 | 10 | 10 |
| | Mean | 97.74 | 5.20 | 56.72 | 28.91 | 18.13 | 9.52 |
| | SD | 0.65 | 1.47 | 3.56 | 3.65 | 2.68 | 1.04 |
| **GILTco1-m-ApoE1** | n | 10 | 10 | 10 | 10 | 10 | 10 |
| | Mean | 97.79 | 5.52 | 54.68 | 30.71 | 19.85 | 9.56 |
| | SD | 0.39 | 1.67 | 3.10 | 2.74 | 1.47 | 1.30 |
| **GILTco1-m-ApoE2** | n | 10 | 10 | 10 | 10 | 10 | 10 |
| | Mean | 98.24 | 3.69 | 55.57 | 31.14 | 19.96 | 9.85 |
| | SD | 0.41 | 0.79 | 4.12 | 4.02 | 3.05 | 1.03 |
| **GILTco1-m-ApoE2-L** | n | 10 | 10 | 10 | 10 | 10 | 10 |
| | Mean | 96.91 | 4.99 | 54.18 | 31.22 | 19.57 | 10.24 |
| | SD | 1.25 | 1.71 | 2.31 | 2.45 | 1.94 | 0.85 |
